# Transposable element disruption of a second thyroglobulin-like gene confers Vip3Aa resistance in *Helicoverpa armigera*

**DOI:** 10.64898/2026.04.06.716841

**Authors:** Andreas Bachler, Tom Walsh, Dan Andrews, Michelle Williams, Tek Tay, Karl Gordon, Bill James, Cao (Grace) Fang, Lisi Wang, Yidong Wu, Eric Stone, Amanda Padovan

## Abstract

**Background:** The cotton bollworm *Helicoverpa armigera* is a major global pest controlled by genetically engineered crops expressing Bacillus thuringiensis (Bt) toxins, including Vip3Aa. While Vip3Aa is widely deployed, the genetic basis of resistance remains poorly understood. Previous work identified disruption of a thyroglobulin-like gene (HaVipR1) as one mechanism of resistance, suggesting additional loci may be involved.

**Results:** Using linkage analysis, transcriptomics, long-read sequencing, and CRISPR-Cas9 gene editing, we identify a second thyroglobulin-like gene, HaVipR2, as a novel mediator of Vip3Aa resistance. Resistance in a field-derived *H. armigera* line was shown to be monogenic, recessive, and autosomal, mapping to chromosome 29. Long-read sequencing revealed a ∼16 kb transposable element insertion disrupting HaVipR2, which was undetectable using standard short-read approaches. CRISPR-Cas9 knockout of HaVipR2 conferred >900-fold resistance, confirming its causal role. Comparative analyses show that HaVipR1 and HaVipR2 share conserved domain architecture, indicating that thyroglobulin-domain proteins represent a recurrent target of resistance evolution.

**Conclusions:** Our findings establish thyroglobulin-domain proteins as a new class of Bt resistance genes in Lepidoptera and demonstrate that transposable element insertions can drive adaptive resistance while evading detection by conventional methods. These results highlight the importance of long-read sequencing and accurate genome annotation for resistance monitoring and provide new insights into the molecular basis and evolution of Vip3Aa resistance.

## Background

The cotton bollworm, Helicoverpa *armigera*, is a major global agricultural pest, traditionally managed with chemical pesticides, but more effectively controlled since the late 1990s through genetically engineered crops expressing insecticidal toxins from *Bacillus thuringiensis* (Bt) (1,2). Bt crops, such as cotton and corn, have significantly reduced pesticide use, promoting sustainability and minimizing off-target ecological impacts (3). In Australia and the United States, Bt crops commonly express pyramids of multiple Bt toxins, frequently combining Cry proteins with Vip3A to delay resistance evolution (4). In Australia, nearly all commercial Bt cotton expresses a three-toxin pyramid comprising Cry1Ac, Cry2Ab, and Vip3A.

While Vip3Aa has been used for control of Heliothine pests for more than a decade, progress in identifying genes or pathways involved in resistance has only emerged recently. Bt toxins target specific insect pests by interacting with receptors or pathways in the insect after ingestion (5,6). Resistance to Bt toxins often results from disruptions to the insect’s receptors or pathways which act to activate the toxin (7). The primary organism where Vip3Aa resistance has been investigated is *Spodoptera frugiperda*, partly due to the availability of the ovarian-derived cell line “Sf9.” In Sf9 cells, several candidates have been proposed, though their relevance in whole insect models remains uncertain (8). *In vivo* studies have provided stronger evidence: resistance has been linked to disruption of the transcription factor SfMyb in a laboratory-derived resistant line (9), and to a retrotransposon insertion within the chitin synthase 2 (SfCHS2) gene, which was shown to confer extreme resistance when disrupted (10). Recent CRISPR validations further extended the role of CHS2 in Vip3Aa resistance across multiple lepidopteran species (11,12). In *H. armigera*, the *HaVipR1* gene (LOC110373801) was disrupted in two independent field-derived Vip3Aa-resistant lines, with the disruption in one line involving a large transposable element insertion in the first intron (13). The *HaVipR1* homologue in *S. frugiperda* was disrupted using CRISPR disruption, with the knockout producing very high levels of resistance, suggesting that VipR1 genes may represent a generalisable resistance locus (14). In *Helicoverpa zea*, genome-wide analyses of Vip3Aa-selected field populations have revealed multiple regions under selection, although specific causal genes have yet to be identified (15,16). Together, these findings highlight that Vip3Aa resistance can arise through disruption of diverse loci, with potential roles for CHS2 and VipR1 across species, as has been widely documented for resistance to Cry1 proteins involving multiple, independent genetic mechanisms.

Transposable elements (TEs) have long been recognized as significant drivers of genomic variability and contributors to resistance mechanisms against Bt toxins. The first Bt resistance gene identified was disruption of a Cadherin gene by the 2.2kb “Hel-1” LTR-type TE (17) in *Heliothis virescens*, conferring Cry1Ac resistance, and in two of the above cases for Vip3Aa resistance it was transposable elements which caused the disruption of VipR1 gene function (10,13). The ability of TEs to disrupt gene function through insertions and mobilization within the genome represents a critical avenue by which pests can develop resistance. With continued improvements in sequencing technologies, particularly long-read sequencing, our ability to identify and characterize TEs has significantly advanced, allowing for direct detection of their role in resistance mechanisms and other genomic functions (see the role of TEs reviewed in plants by (18)). Despite these technological advancements, challenges remain. Even with improved long-read methodologies, certain large structural variations, including TE insertions, can be overlooked by traditional mapping and reference-based approaches. *De novo* assembly methods offer a more robust alternative, reducing the impact of reference bias and enabling a more comprehensive detection of genomic variation (19).

In this study, we explore a novel field-derived Vip3Aa-resistant line of *Helicoverpa armigera* that is non-allelic to previously identified resistant lines. Through sex-based crosses and short-read sequencing, we pinpoint the chromosome and region associated with this resistance. Subsequently, we employed long-read sequencing to characterize a TE insertion as a novel mechanism of resistance. Our findings underscore the limitations of short-read sequencing in identifying significant genetic variations and demonstrate how these limitations can obscure identification of critical resistance genes. This work not only highlights the role of TEs in conferring resistance, demonstrating a genetically and mechanistically distinct basis of Vip3Aa resistance from previously identified loci, but also emphasizes the necessity of employing advanced sequencing technologies to fully understand and manage resistance in agricultural pests globally, as Bt crops with Vip3Aa continue to be developed and implemented.

## Results

### Linkage analysis identifies Vip3Aa resistance region and candidate resistance gene

From 35 single-pair crosses, 34 yielded valid results (one was excluded due to low survival in the control population). Among these, 25 crosses were Male-Informative Crosses (MIC) at both the F_1_ and F_2_ stages, five were Female-Informative Crosses (FIC) at both the F_1_ and F_2_, and four were a mixed setup of FIC at F_1_ and MIC at F_2_ (see **Supplementary Figure S1** for crossing diagram and **Supplementary Table S1** for sequence information obtained for all samples). MICs were prioritized for their potential to improve resolution through chromosomal crossover, while FICs served to confirm the chromosome associated with resistance. The direction of the cross did not affect survival ratios (**Figure 1 A**), indicating resistance was not linked to the sex chromosome. The average survival rate on treated plates was 41.7%, compared to 90.5% on control plates. A chi-square test for monogenic inheritance (p = 0.29) supported a monogenic resistance model for Vip3Aa resistance.

**Figure 1.**
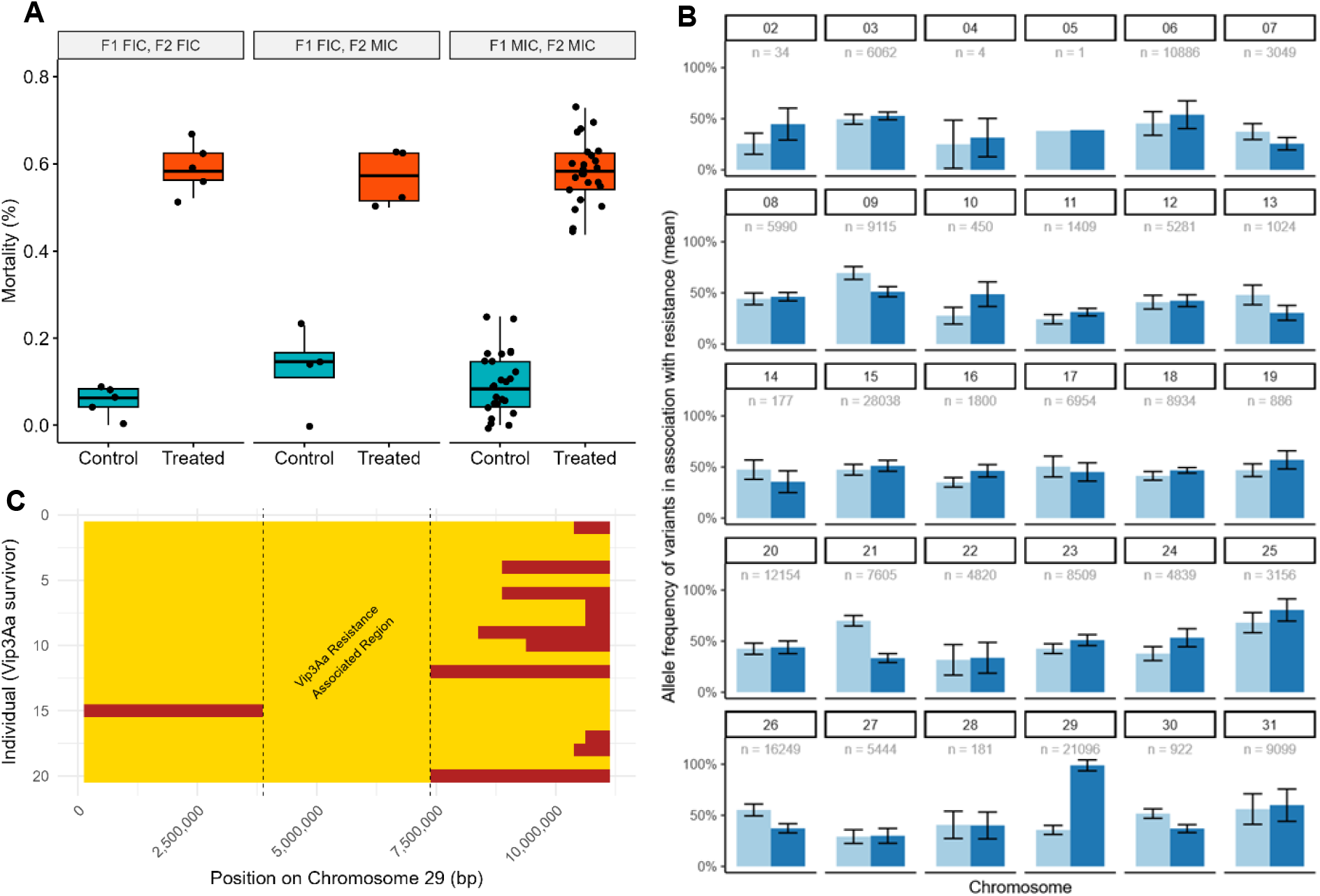
Genetic mapping of Vip3Aa resistance in *H. armigera*. (A) Survival of progeny from single-pair crosses exposed to Vip3Aa. 34 of 35 crosses produced viable data, including 25 male-informative crosses (MIC) and nine female-informative or mixed crosses (FIC). A chi-square test supported a monogenic resistance model. (B) Genome-wide analysis of maternally derived (i.e. Vip3Aa resistance linked) marker SNPs in control (light blue) and treated (dark blue) populations. Among 368,336 informative variants, only chromosome 29 showed near fixation of resistance alleles in treated individuals, while controls remained heterozygous. (C) Fine-scale mapping of chromosome 29 using MIC data. Median allele frequencies across 250-kbp windows revealed a single, contiguous region (3.9–7.4 Mb) homozygous in all resistant progeny, delineating the Vip3Aa resistance locus. Dark-red = heterozygous regions; yellow = homozygous regions.

Short-read sequencing was used to assess inheritance in F_0_, F_1_, and F_2_ individuals from a Female-Informative Cross (FIC). High-quality SNPs informative of the resistance genotype were identified, resulting in 368,336 marker variants across all autosomes. These were genotyped in treated and control populations (**Figure 1 B**). Chromosome 29 was the only chromosome where allele frequencies approached fixation in treated populations, while control populations remained around 50% heterozygosity. Fine-scale mapping using MIC data further resolved the resistance region. Bi-allelic SNPs were binned in 250-kb segments, revealing a single contiguous block from 3.75–7.25 Mb on chromosome 29 in which all variants were homozygous in Vip3Aa survivors but remained heterozygous in controls (**Figure 1 C**). This interval was therefore defined as the resistance-associated region.

Assessment of the chromosome 29 resistance-associated region revealed one gene with similarity to the recently described Vip3Aa resistance gene HaVipR1. This gene (LOC126056805) contains two thyroglobulin type-1 repeat domains and emerged as the sole gene in the region with functional characteristics with any known Vip3Aa resistance factors. We therefore designate this gene HaVipR2. To determine whether HaVipR2 could represent a paralogous Vip3Aa resistance gene, we compared its sequence, structure, and functional annotations with HaVipR1. HaVipR1 is located on chromosome 27 and encodes a 420-aa protein across nine exons, while HaVipR2 lies on chromosome 29 and encodes a 330-aa protein across seven exons (using chromosome numbering from the HaSCD2 reference GCF_023701775.1). Despite low primary-sequence identity (24% amino-acid identity), both proteins contain the same characteristic domain architecture: two thyroglobulin type-1 repeats (PF00086), the Secreted Modular Calcium-Binding Protein family (PTHR12352), and identical GO assignments (BP: cell-matrix adhesion (GO:0007160) and CC: extracellular space (GO:0005615); basement membrane (GO:0005604)). InterProScan assigned both as part of the “Developmental regulatory and protease inhibitor” family (IPR051950). AlphaFold3 structural models showed high-confidence predictions for both proteins (after excluding N-terminal signal peptide regions). Superposition of the trimmed structures indicated strong similarity, with an RMSD of 3.301 Å. Across the *H. armigera* genome, only three additional genes contain any thyroglobulin-type domains, and none share the specific domain architecture or family assignments characteristic of HaVipR1 and HaVipR2 (**Error! Reference source not found**.).

Taken together, the linkage mapping and comparative analyses identify HaVipR2 as the primary candidate resistance gene within the chromosome 29 locus.

**Table 1.** Other genes containing thyroglobulin-1 domains in the *H. armigera* genome. Genes were identified based on presence of thyroglobulin-1 domain (pfam: PF00086; InterProScan: IPR000716) of protein sequences using InterProScan v5.

| Gene ID & chromosome | Gene Name | Comment |
| --- | --- | --- |
| LOC110374505<br>Z Chromosome | SPARC-related modular calcium-binding protein 2 | 2 thyroglobulin-1 domains identified. Additional domains include SPARC/testican calcium binding domain, Kazal/serine protease inhibitor domain and EF-hand-domain |
| LOC110383208<br>Chromosome 9 | Proteoglycan Cow | 1 thyroglobulin-1 domain identified. Additional domains include SPARC/testican calcium binding domain, Kazal/serine protease inhibitor domain and EF-hand domain |
| LOC110382892<br>Chromosome 25 | Extracellular matrix protein A | 6 thyroglobulin-1 domains identified. Additional domains include trypsin inhibitors, hirudin/antistatin and a secreted calcium binding domain. |

### Gene expression profiling of Vip3Aa resistant and susceptible lines

Transcriptomic analysis was conducted on resistant and susceptible lines to attempt to identify mechanisms underlying resistance in the identified region. Differential Gene Expression (DGE) analysis compared a susceptible line (“GR”) without Vip3A exposure to a resistant line (“17-294”) under both Vip3A-treated and untreated conditions. Mid-gut samples were pooled from five individuals per group, with three biological replicates per condition.

Principal component analysis (PCA) revealed clear separation between the susceptible and resistant lines, with the first principal component capturing most of the variation (**Figure 2 A**). Although one replicate of the susceptible line (“GRxUnt_2”) showed slight divergence, it remained distinct from the resistant lines. There was little impact of Vip3Aa treatment seen in the resistant line, with overall similar expression profiles seen in the resistant line with or without Vip3Aa treatment, corroborated by heatmap clustering (**Figure 2 B**).

**Figure 2.**
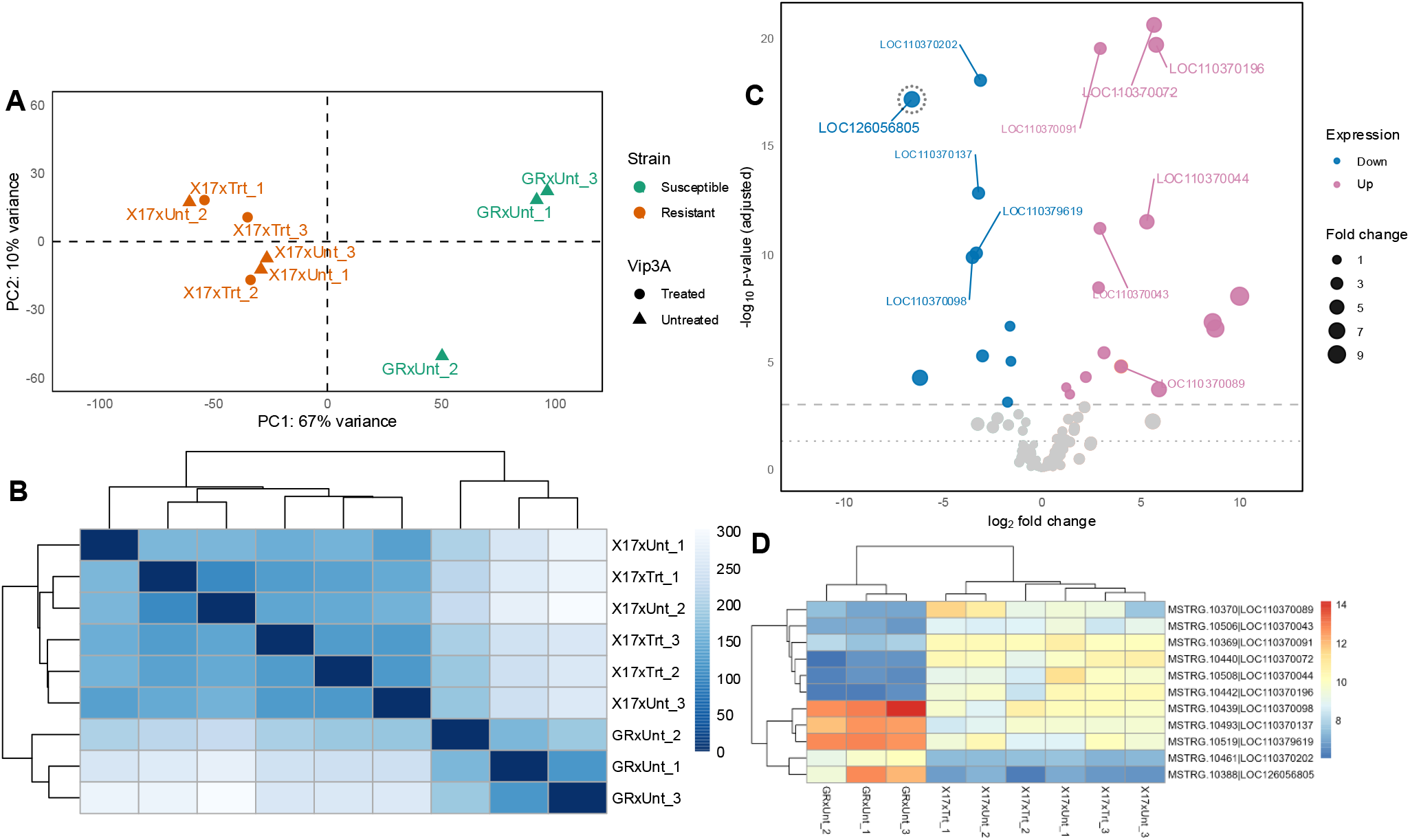
Transcriptomic analysis of Vip3Aa resistance *in H. armigera*. (A) Principal component analysis (PCA) of variance-stabilized gene expression from pooled midgut tissue of susceptible (“GR”) and resistant (“X17”) larvae, with (“Trt”) and without (“Unt”) Vip3Aa treatment. Biological replicates (n = 3 per condition; 5 pooled midguts each) separated primarily by genetic background, with treatment having little global effect. (B) Euclidean distance heatmap of gene expression of the same samples confirms clear clustering by resistance phenotype rather than treatment. (C) Volcano plot of differential gene expression (DGE) within the chromosome 29 resistance locus identifies 25 significantly altered genes (adjusted *p* < 0.001) between resistant and susceptible lines. Dashed and dotted lines indicate p-value thresholds of 0.001 and 0.01, respectively. Notably, a thyroglobulin-like gene (LOC126056805) is strongly down-regulated in resistant samples. (D) Heatmap of the top 10 differentially expressed genes in the resistance region shows consistent expression shifts distinguishing resistant from susceptible lines under both treated and untreated conditions.

Assessing DGE overall between just the susceptible (“GR”) and resistant (“X17”) lines (and excluding the impact of Vip3Aa treatment) revealed thousands of differentially expressed genes (n = 3024, with adjusted p-value < 0.01), indicating a substantial difference in the transcriptional profiles of the resistant and susceptible strains. These were filtered to assess genes previously identified as being involved in Vip3A resistance in *Spodoptera frugiperda*. The raw differential expression values of these candidate genes in the resistant line are provided in **Supplementary Table S2** and the transformed values for all samples shown in **Supplementary Figure S2**. None of the candidate genes were found in the region associating with resistance identified from the crossing experiment. There are two genes found to be differentially expressed that have been previously found to be involved in Vip3A resistance in *S. frugiperda*: “fibroblast growth factor receptor homolog 1” (FGFR / LOC110373728) and “autophagy protein 5” (ATG5 / LOC110374447)) however, as mentioned, these were not found in the region associating with resistance.

Within the resistance-associated region on chromosome 29, 25 genes were significantly differentially expressed between resistant and susceptible samples (adjusted *p* < 0.001) (**Figure 2 C**). Among these, the “thyroglobulin-like gene” (LOC126056805, referred to here as “HaVipR2”) exhibited a 6-fold decrease in expression. The top ten most differentially expressed genes are listed in **Supplementary Table S3**. The thyroglobulin-like gene was significantly down-regulated in resistant samples compared to susceptible ones (**Figure 2 D**).

Vip3Aa treatment in the resistant line led to a small number of differentially expressed genes (n=24, adjusted p < 0.01), none of which were located within the resistance associating region on chromosome 29. Among these, two apoptosis-related genes, ‘croquemort’ (LOC110382852) and ‘caspase-8’ (LOC110376609), were significantly up-regulated in Vip3Aa-treated resistant lines (**Supplementary Table S4**).

Transcriptomic assessment revealed clear expression differences between resistant and susceptible lines, with Vip3Aa exposure itself causing little additional transcriptional change, consistent with a high level of Vip3Aa resistance. Within the resistance-associated region on chromosome 29, several differentially expressed candidate genes were identified, but none corresponded to any of the previously described Vip3Aa resistance genes from *S. frugiperda*.

### CRISPR-Cas9 disruption of *HaVipR2* confirms its role in Vip3Aa resistance

#### Establishment of the knockout strain of HaVipR2

A total of 1,220 fresh eggs were individually injected with a 1-nanoliter mix of Cas9 protein and two *HaVipR2* sgRNAs. Of these, 17.05% (208/1,220) hatched. Among the 208 neonates, 60.58% (126/208) developed into adults (G_0_). Thirty G_0_ moths were randomly chosen for single-pair crosses with SCD moths to produce the next generation (G_1_). Fourteen out of the 30 single-pair families produced fertile progeny. Genomic DNA from pools of ten second instar larvae from each of these 14 families was screened for indel mutations. Nine families exhibited at least one Cas9-induced indel mutation, with one family (#13) harbouring a 95-bp deletion in exon 1 of *HaVipR2*.

Sixty G1 moths from family #13 were randomly selected for genotyping at exon 1 using a non-destructive approach. Among them, 30 moths (14 ♀ and 16 ♂) were identified as heterozygous for the 95-bp deletion and were mass-crossed to produce G2 progeny. Of the 70 G2 moths genotyped, 16 were homozygous for the 95-bp deletion. These homozygotes were pooled and mass-crossed to establish a homozygous strain, designated as *HaVipR2*-KO. The 95-bp deletion is predicted to introduce a premature stop codon, leading to a truncated, non-functional protein.

#### Response to Bt proteins of the susceptible and knockout strains

The LC50 of Vip3Aa for the susceptible SCD strain was determined to be 0.142 μg/cm^2^ (95% CI: 0.054 - 0.358). In contrast, the *HaVipR2*-KO strain exhibited only 20.8% (5/24) survival when exposed to 128 μg/cm^2^ of Vip3Aa (maximum tested), indicating a resistance level more than 900-fold higher than that of the SCD strain.

#### Genetic linkage between knockout of HaVipR2 and resistance to Vip3Aa

A backcrossing strategy was employed to confirm the causal relationship between *HaVipR2* knockout and Vip3Aa resistance. Fifteen male moths from the *HaVipR2*-KO strain were mass-crossed with 15 female SCD moths, and vice versa. The resulting F1 offspring were exposed to a diagnostic concentration of Vip3Aa (4 μg/cm^2^), resulting in 99.3% (286/288) mortality, indicating nearly complete recessive resistance. Fifteen F1 males were then mass-crossed with 15 *HaVipR2*-KO females to produce backcross progeny (BC1). Upon exposure of 144 BC1 neonates to Vip3Aa, 45.14% (65/144) survived. Genotyping of 30 survivors revealed all were homozygous for the 95-bp deletion, compared to 17 homozygotes and 13 heterozygotes among the 30 untreated larvae. These results strongly link the *HaVipR2* knockout with Vip3Aa resistance, as confirmed by Fisher’s exact test (*p* < 0.01).

#### Cophylogenetic analysis of *HaVipR1* and *HaVipR2*

Phylogenetic reconstruction of *HaVipR1* and *HaVipR2* across representative Lepidoptera revealed largely congruent topologies but with notable differences at key nodes (**Figure *3***). In both trees, sequences from Noctuoidea and Tortricoidea consistently clustered together with strong ultrafast bootstrap support (>90%), indicating that both *HaVipR* paralogs have maintained conserved evolutionary trajectories within these superfamilies.

**Figure 3.**
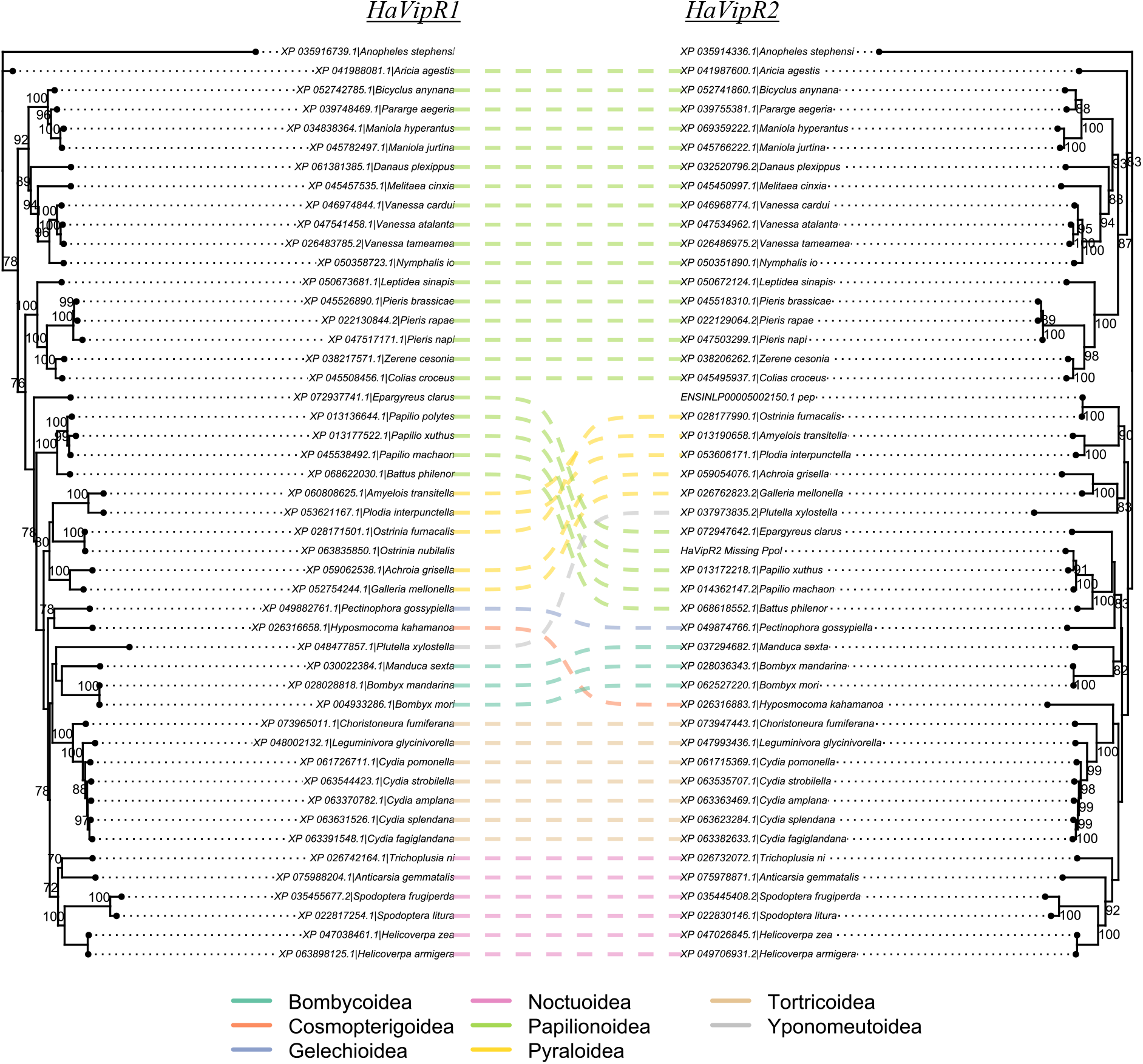
Phylogenetic relationships of HaVipR1 and HaVipR2 genes. Cophylogeny of *HaVipR1* amino acid sequences from representative Lepidoptera. Sequences were retrieved from RefSeq Lepidoptera genomes, aligned with MAFFT, and phylogenies inferred with IQ-TREE using the ModelFinder option. Anopheles stephensi was specified as the outgroup. Links between clades are colored by superfamily. Node values indicate ultrafast bootstrap support; only nodes with support >70% are shown.

The placement of single representatives from Cosmopterigoidea (*Hyposmocoma kahamanoa*), Yponomeutoidea (*Plutella xylostella*), and Gelechioidea (*Pectinophora gossypiella*) differed slightly between the *HaVipR1* and *HaVipR2* phylogenies; however, these nodes were supported only weakly (<70%), suggesting that their relationships remain uncertain.

A difference was observed in the position of Pyraloidea. In the *HaVipR1* phylogeny, Pyraloidea formed a cohesive clade sister to Papilionoidea, whereas in the *HaVipR2* tree, Pyraloidea was displaced, effectively splitting Papilionoidea into two groups. This topological discordance likely reflects either lineage-specific divergence in *HaVipR2* or reduced phylogenetic signal in this clade.

Overall, the co-phylogeny demonstrates that *HaVipR1* and *HaVipR2* have followed broadly parallel evolutionary histories across Lepidoptera, while also revealing clade-specific differences that may underlie functional diversification of these paralogs.

### Mis-annotation of *HaVipR2*

The sequence of *HaVipR2* appears to be less reliably annotated than *HaVipR1*. It was first identified as missing from the RefSeq annotation of the initial *H. armigera* genome (Harm1.0, GCF_002156985.1), although the sequence for the gene was present in the genome and RNA-Seq indicated expression at the location. This gene was present in the annotation generated by Helicoverpa Genome Consortium for the same genome (GCA_002156985.1; HaOG205528).

Using the HaVipR1 and HaVipR2 protein sequences as a query of the RefSeq database with blastp (filtering to keep only species with a match E-value < 1^e-100^) there are two species which have a match for *HaVipR1* but do not appear to have one for *HaVipR2* in their protein sequences (derived from the genome annotation). These two organisms are *Papilio polytes* (“Common Mormon”; GCF_000836215.1) and *Ostrinia nubilalis* (“European corn borer “; GCF_963855985.1). Re-annotation of the genome sequences using metaeuk with *HaVipR1* and *HaVipR2* indicates that in both cases the gene appears present in the genome but is missing from the protein annotation. In the case of *P. polytes* the gene is mis-annotated as a ‘pseudogene’ (LOC106104612) and is therefore not converted to a protein sequence and not included in the blastp search. For *O. nubilalis*, the gene is simply missing from the annotation, although it is present in the Ensembl annotation for this organism (Gene: ENSINLG00005001987.1; Protein: ENSINLP00005002150.1).

### Transposable element disrupting *HaVipR2* is hidden from short-read based analysis

The transcriptomic evidence and CRISPR-Cas9 disruption of *HaVipR2* indicate that down-regulation or disruption of this gene confers high levels of Vip3Aa resistance. Analysis of short-read whole-genome sequence (WGS) data from the resistant individuals was initially unsuccessful in identifying a disruptive variant in the *HaVipR2* gene using a variety of approaches, including structural variant detection and variant annotation with a variety of bioinformatic tools (described in Supplementary Methods Section 2).

An analysis of structural variants utilizing *de novo* assemblies of long-read WGS data from the resistant individuals indicated that there was a large insertion (∼16,000bp) event present in the coding region of exon 5 of the *HaVipR2* gene. This was identified as a transposable element with two large (∼4000bp) inverted repeat sequences on either side of the element and 9-bp target-site duplication (TSD) present at the site of insertion (**Figure 4**). The large flanking repeats and the TSD led to overlapped mapping from both short and long-read alignments, with structural variant callers unable to correctly identify the event as an insertion, sometimes mis-classifying it as a ‘break-point’ event, due to partial mapping to other regions in the *H. armigera* genome.

**Figure 4.**
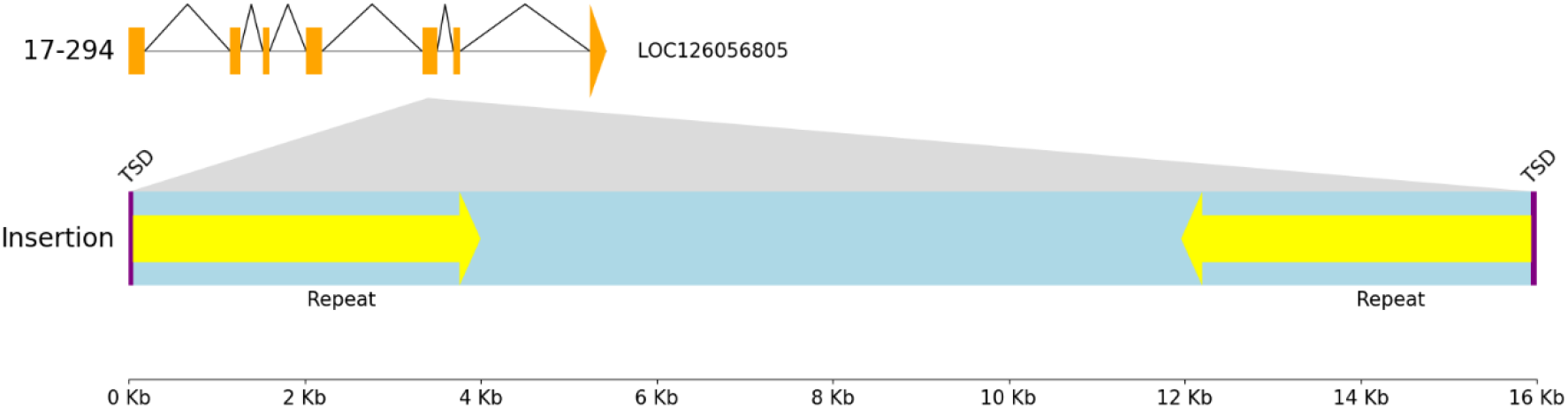
Structure of the large insertion identified from the *de novo* assembly of a resistant individual. The assembled sequence was annotated to identify the gene structure of *HaVipR2*, and repetitive elements were detected with RepeatFinder in Geneious Prime. The insertion is flanked by a 9-bp target site duplication (TSD; GGA-ACA-AAT). Immediately adjacent to these TSD sequences are two large inverted repeats of 3,973 bp with 97.1% identity. No open reading frame longer than 200 bp was detected within the insertion region.

The long-read *de novo* assembly from the resistant individual with the disruptive transposable element was used as a reference to align RNA-Seq from the resistant and susceptible individuals. Aberrant splicing was identified for this gene in the resistant individuals, confirming that the transposable element disrupts normal gene function in the resistant individuals (See **Supplementary Figure S5**).

The large transposable element insertion region was assessed for presence in other *Helicoverpa* genomes. The results from blastn of the large insertion element indicated that there were 9 distinct regions in *Helicoverpa armigera armigera* (HaSCD2/ GCF_023701775.1) which had >99% query cover and ∼90% identity and 14 distinct regions in *Helicoverpa zea* (ASM2234304v1/ GCA_022343045.1) which also had >99% query cover and ∼90% identity. None were identified on chromosome 29 which is where resistance has been identified in this instance.

## Discussion

This study identifies a second thyroglobulin-like gene, *HaVipR2*, as a novel gene involved in Vip3Aa resistance in *Helicoverpa armigera*. Using bi-phasic linkage analysis, transcriptomics, long-read sequencing, and CRISPR validation, we demonstrate that resistance in the field-derived line ‘17-294’ is monogenic, recessive, and autosomal, and results from the insertion of a large transposable element that disrupts the coding sequence of *HaVipR2*. Together with our earlier identification of *HaVipR1*, these findings establish thyroglobulin-domain proteins as a previously unrecognized class of Bt resistance genes in Lepidoptera. Unlike most Cry Bt resistance mechanisms, which involve alterations in toxin receptors such as cadherins, ABC transporters, or alkaline phosphatases (6,20), disruption of *HaVipR1* and *HaVipR2* potentially highlight an alternative pathway for resistance.

Initial identification of the disruption in *HaVipR2* was complicated by the repetitive nature of the transposable element which was inserted into the coding sequence of the *HaVipR2* gene. Our bi-phasic cross design proved highly effective for localizing the resistance locus, with female-informative crosses identifying the resistance chromosome and recombinant male-informative crosses narrowing the interval to chromosome 29. However, attempts at using short-read mapping methods failed to correctly detect the large transposable element insertion in the *HaVipR2* gene, either missing or misclassifying the variant, despite clear dysregulation present for the gene from transcriptomic analysis. Only a long-read *de novo* assembly approach correctly identified the causal insertion (21). With the majority of Bt resistance arising from disruptions of gene function this highlights the known problems associated with reliance on reference-based mapping approaches, which can overlook complex structural variants, particularly large TE insertions (22). Monitoring for specific resistance alleles continue to be challenging with marker or short-read based methods as these are often limited to identification of specific variants, while as we demonstrate here, long-read based methods appear to provide a more comprehensive approach for discovering such variants and should be considered, where appropriate, for future resistance surveillance programs.

The disruption of *HaVipR2* mirrors that of its paralogue *HaVipR1*, which is present on a different chromosome but encodes a protein of similar function and structure. The repeated involvement of this gene family strongly suggests that disruption of thyroglobulin-domain genes represents a generalizable mechanism of Vip3Aa resistance in *H. armigera*. Consistent with this, recent CRISPR-mediated disruption of *SfVipR1* in *Spodoptera frugiperda* conferred high levels of Vip3Aa resistance (14), suggesting that VipR-family genes may represent a broader resistance pathway across Lepidoptera. In *H. zea*, genome-wide selection scans have revealed multiple genomic regions associated with Vip3Aa resistance, though causal genes have not yet been identified (15,16). Assessment for the identified regions for HaVipR genes may provide indication of selection of resistance in these areas.

Mechanistically, the involvement of two genes on distinct chromosomes, but with similar functional annotations, may support a model in which Vip3Aa resistance arises from altered regulation of cellular repair rather than direct modification of toxin binding. Both *HaVipR1* and *HaVipR2* have Thyr1 domains, which are associated with cysteine protease inhibitors (23,24). Loss of these inhibitors may lead to elevated cysteine protease activity, enhancing membrane repair and shifting cells toward survival pathways such as autophagy rather than apoptosis. This interpretation is supported by the consistent higher expression of ATG5 in the resistant line in comparison to the susceptible line, and specifically from altered expression of apoptosis-related genes in resistant larvae exposed to Vip3Aa, and parallels mammalian studies where cysteine proteases released from lysosomes play central roles in plasma membrane repair (25). There is some evidence from studies of Vip3Aa in *Sf9* cells that indicates increase in autophagy may improve resistance (26) and also that Vip3Aa is potentially present in lysosomes (27). While this provides some support for the action of Vip3Aa in *Sf9* cells, previous investigation of *HaVipR1* expression in Sf9 cells indicated it was expressed at incredibly low levels, making it challenging to consider a situation where a disruption would lead to resistance, since the expression level is already so low. However, it may be that the low expression is part of the Sf9 cell profile, and despite this low expression, *HaVipR2* functions in a similar way in the Sf9 cell line as it does in the insect midgut. Another limitation of the current model is its inability to explain the absence of cross-resistance to Cry toxins, which also act through pore formation, and so presumably changes in repair should lead to increased resistance to Cry toxins in addition to Vip3Aa (28). This suggests that thyroglobulin-mediated repair may act selectively on Vip3Aa-induced pores, or that additional gene-specific or structural differences mediate the specificity of the response.

Despite this emerging model, functional and structural information for the HaVipR proteins remains poor. Further characterisation of these genes could indicate other roles in the mid-gut, which may lead to identification of resistance pathways more directly linked to Vip3Aa activation or binding, rather than shifts in cellular repair. Structural analysis of the HaVipR genes and biochemical assays will be essential to determine whether HaVipR proteins interact directly or indirectly with Vip3Aa. Comparative genomics also reveals that HaVipR genes can be misannotated either as pseudogenes or missed altogether in other Lepidoptera, emphasizing the need for careful re-annotation to fully understand their distribution and function.

Taken together, these results highlight a broader challenge in detecting the genomic basis of adaptive traits, which is that complex structural variants, particularly those mediated by transposable elements, remain difficult to identify with short-read data and standard reference-based workflows. In this study, a ∼16 kb insertion within the HaVipR2 gene was entirely missed or misclassified by multiple widely used short-read callers (see Methods below, and Supplementary Section 2), despite clear functional disruption observed at the transcriptomic level. Only long-read sequencing combined with a *de novo* assembly-based SV method (SVIM-asm) resolved the insertion properly. The challenge of accurately assessing the impact of TE elements for Bt resistance have long been noted in Bt resistance research, beginning with the Hel-1 transposable element insertion that disrupted cadherin (17), and more recently in TE-mediated disruptions of HaVipR1 and SfCHS2. Our findings therefore reinforce that TE insertions represent a recurrent, and likely under-detected, source of resistance alleles in Lepidoptera. The discovery that HaVipR2, the paralogue of HaVipR1, is independently disrupted by a large TE insertion indicates that thyroglobulin-domain genes form a recurrent target of Vip3Aa resistance evolution in H. armigera. The repeated involvement of this gene family across independent resistant lines suggests that loss of thyroglobulin-domain function may represent a common route by which Vip3Aa resistance can arise. More generally, the difficulty of detecting this insertion with short-read methods underscores the limitations of conventional surveillance approaches for resistance alleles, which often rely on SNP markers or targeted assays that cannot capture complex SVs. As long-read sequencing becomes more widely accessible, integrating assembly-based SV discovery with careful gene annotation will be essential for accurately characterizing resistance mechanisms, monitoring their spread, and understanding how TE activity contributes to rapid evolutionary change in pest species.

## Materials and Methods

### Insect lines and rearing

Susceptible *Helicoverpa armigera* strain “GR” and Vip3Aa-resistant strain “17-294” (first isolated by Chakroun et al., 2016) were maintained at 25 ± 1 °C, 50 ± 10% relative humidity, and a 14:10 h light:dark cycle at the CSIRO Black Mountain Laboratories (Canberra, Australia). All colonies were reared on solid artificial diet prepared according to (29). Adults were maintained in 750 mL perforated polypropylene containers (Décor FoodFresh #051900) and provided with fortified honey solution (29). Eggs were collected on cloth oviposition substrates and stored at 8 °C until hatching.

Single-pair crosses between resistant (17-294) and susceptible (GR) individuals were established to produce F_1_ progeny. F_1_ pupae were sexed and heterozygous individuals backcrossed to resistant moths to generate F_2_ families. Crosses were categorized as Female-Informative Crosses (FICs) when the heterozygous parent was female, enabling chromosome-level linkage detection due to the absence of recombination in female Lepidoptera. Crosses in which the heterozygous parent was male were designated Male-Informative Crosses (MICs) and were used for fine-scale recombination-based mapping of the resistance locus (30).

### Vip3Aa bioassays and F_2_ selection

Phenotyping of F_2_ progeny was performed using surface-overlay bioassays in 48-well plates (0.2 cm^2^ surface area/well). Solid diet was overlaid with 1.69 µg/cm^2^ Vip3Aa, previously determined to cause 100% mortality in the susceptible GR line and consistent with high mortality in susceptible *H. zea* (31). Control plates received an equivalent volume of H_2_O overlay.

For each cross, 48 neonates were placed onto treated wells and 48 onto paired control wells. Larvae were scored after 5 days, using criteria adapted from Chakroun et al. (2016), with individuals were classified as dead (including moribund/immobile) or alive. Plates were then frozen to euthanize larvae. All survivors from treated and control plates were collected for DNA extraction.

Two representative crosses - one FIC and one MIC - were selected for DNA sequencing. All survivors from these crosses were processed for short-read sequencing, and the four parental individuals from the MIC (F_0_ and F_1_) were used for long-read DNA extraction. One additional member of the resistant line 17-294 was also chosen for follow-up sequencing to confirm the large insertion was present.

### Statistical analysis of monogenic inheritance

The inheritance of Vip3Aa resistance was tested for consistency with a monogenic model following (32). For each F_2_ backcross, the expected proportion of resistant progeny under a single-locus RR × RS model is 50%. A chi-square goodness-of-fit test was applied using:

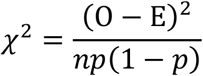

where O is the number of observed survivors, E is the expected number (np), n is total larvae assayed, and p = 0.5. The null hypothesis of monogenic inheritance was rejected when P < 0.05 (degrees of freedom = 1).

### DNA extraction and sequencing

#### Short-read sequencing

Genomic DNA was extracted from whole larvae or dissected tissue using the Qiagen DNeasy 96 Blood & Tissue Kit (Qiagen, Hilden, Germany). DNA quality was verified by NanoDrop and Qubit fluorometry. Libraries were prepared using the Illumina DNA Prep-M kit (300-cycle format) and sequenced (2 × 150 bp) on an Illumina NovaSeq S4 lane at the Australian Genome Research Facility (AGRF).

#### Long-read sequencing

High-molecular-weight (HMW) DNA was extracted from thorax tissue using the Circulomics Nanobind HMW kit. DNA integrity was assessed using the Promega QuantiFluor dsDNA system. Libraries were prepared using ONT’s Ligation Sequencing Kit (Kit 14 chemistry, SQK-LSK114) on the PromethION P2 platform (R10.4.1 flow cells, FLO-PRO114M). The first 4 samples (all MIC cross individuals) were sequenced at the Biomolecular Resource Facility (BRF), Canberra, while the 2 follow-up samples were sequenced at the Australian Genome Research Facility (BRF). High-accuracy base-calling was performed, and reads were concatenated for downstream analysis.

### Long-read *de novo* assembly

#### The susceptible “GR” reference genome

High-molecular-weight DNA was extracted from 50mg thorax tissue of a single moth from the susceptible GR line using the Circulomics Nanobind HMW kit. Long-read libraries were prepared and sequenced on the ONT PromethION platform with raw reads being error-corrected and assembled using Canu (v2.2.0; (33)) with the genome size parameter set to 380 Mb. The resulting assembly was polished with three rounds of Pilon ((34)) using the matched Illumina short-read data. Haplotigs and redundant sequences were purged using Purge_Haplotigs (v.1.1.2; (35)). Hi-C scaffolding was then performed with YaHS (v1.2.2; (36)), generating 31 major chromosomal scaffolds. Chromosome numbering was determined based on alignment to the HaSCD2 reference genome using Mashmap (v.2.0.1; (37)). The mitochondrion was assembled *de novo* from short-read data using Mitofinder (v.1.4.1; (38)) and concatenated to the nuclear genome. Putative W-chromosome scaffolds were identified by analyzing female (ZW) versus male (ZZ) short-read coverage; unplaced scaffolds >100 kb in length with >15% 1000bp windows exclusively mapped in ZW samples were concatenated into a pseudo-W chromosome. Assembly completeness was assessed with BUSCO (v5.2.2) using the Lepidoptera_odb10 database, and repeat annotation was performed using RepeatModeler (v.2.0.2; (39)) and RepeatMasker (v.4.1.2). Genome annotation was conducted using Liftoff (v.1.6.3; (40)) with the annotated HaSCD2 genome (GCF_023701775.1) as the reference with default parameters. Functional annotation was conducted with InterProScan (v5; (41)). The final genome sequences was submitted to GenBank.

#### All other individuals

Other F_0_/F_1_ individuals, and the follow-up 17-294 resistant individual, were assembled *de novo* using Flye (v.29.3; (42)) with default settings, followed by three rounds of polishing with matched short reads using Pilon.

### Identification of the resistance-associated genomic region

Reads from all F_0_, F_1_, and F_2_ individuals were trimmed with fastp and aligned to the *H. armigera* “GR” reference genome using BWA-MEM2 (v2.2.1;(43)). Pairing information was corrected (samtools fixmate (v1.18; (44))), duplicates removed (Picard MarkDuplicates), and alignments sorted and indexed with samtools.

Variants were jointly called using FreeBayes (v1.3.7; (Garrison & Marth, 2012) with adjusted parameters (*--use-best-n-alleles 16 --min-alternate-count 2 --min-coverage 5 --no-population-priors --report-genotype-likelihood-max*). Variants were filtered using bcftools (QUAL > 20; depth ≥ 5; no nearby INDELs). Average read depth across all individuals was approximately 20×.

#### FIC mapping

To identify the chromosome carrying the resistance allele, filtered SNPs were selected according to the expected resistance-inheritance pattern (resistant parent: hom-alt; susceptible parent: hom-ref; F_1_: het). Alleles in treated and control F_2_ groups were compared for inheritance of these resistance-associated SNPs. The chromosome showing fixation of the resistant alleles in treated offspring was designated as carrying the resistance locus.

#### MIC fine-mapping

Variants on the identified chromosome were further filtered to high-quality bi-allelic SNPs supported by ≥10 reads. Allele frequencies in treated and control F_2_s were summarized in 250 kb sliding windows. Regions with the majority of SNPs as homozygous exclusively in treated individuals were defined as resistance-associated intervals.

### Transcriptome sequencing and differential expression analysis

Third-instar larvae from GR and 17-294 strains were reared under standard conditions described above. For transcriptome profiling, 64 individuals per condition were placed individually into wells containing either untreated diet (GR and 17-294) or Vip3Aa-overlay treated diet (17-294 only), using the same toxin concentration as in the selection experiments. Larvae were transferred to fresh plates every 2–3 days to maintain exposure.

Upon reaching fifth instar, midguts from five individuals were pooled to form one biological replicate (three replicates per condition). RNA was extracted using the Zymo Quick-RNA Tissue/Insect MicroPrep Kit, and quality assessed with Agilent TapeStation. Libraries were prepared and sequenced on an Illumina NovaSeq S1 lane at AGRF.

Reads were mapped to the “GR” reference using Hisat2 (v2.2.1; (46)). Transcript abundance was quantified using StringTie (v3.0.1; (47))and merged to form a unified gene-level count matrix. Differential expression was performed in DESeq2 (v3.22; (48)) with variance-stabilizing transformation for visualization (pHeatmap). Analyses focused on (i) genome-wide differences between resistant and susceptible lines, (ii) genes located within the resistance-associated interval and (iii) differential gene expression patterns between the treated and untreated resistant lines.

### CRISPR-Cas9 disruption of *HaVipR2* confirms its role in Vip3Aa resistance

#### Insect strains

The wild-type strain SCD that was originally collected from Côte D’Ivoire (Ivory Coast, Africa) in the 1970s was kindly provided by Bayer Crop Science in 2001 (49). This strain has been maintained in the laboratory without exposure to insecticides or Bt toxins for over 40 years, and it is susceptible to both Bt toxins and chemical insecticides (49). *HaVipR2* gene (LOC126056805) of the SCD strain was knocked out with CRISPR/Cas9 genome editing to produce a knockout strain, *HaVipR2*-KO, which is homozygous for a 95-bp deletion in exon 1 of *HaVipR2*.

#### Vip3Aa protein and bioassay

The Vip3Aa used in this study was provided by the Institute of Plant Protection, Chinese Academy of Agricultural Sciences (CAAS), Beijing, China.

Toxicity of Vip3Aa to SCD and the knockout strain were determined with diet overlay bioassays. Vip3Aa solutions were prepared by diluting the stock suspensions with a 0.01 M, pH 7.4, phosphate buffer solution (PBS). Liquid artificial diet (1000 µl) was dispensed into each well (surface area = 2 cm^2^) of a 24-well plate. After the diet cooled and solidified, 100 µl of Vip3Aa protein solution was applied evenly to the diet surface in each well. A single unfed neonate (24 h old) was put in each well after the Bt protein solution was dried at room temperature, and mortality was recorded after 7 days. Larvae were considered as dead if they died or weighed less than 5 mg at the end of bioassays.

#### Design and preparation of a single guide RNA (sgRNA) targeting exon 1 of HaVipR2

Two sgRNA target sequences (sgRNA-1: 5’-TGTAAGCCTACAGTGTCAGATGG-3’; sgRNA-2: 5’-TATGGGTGCTGTCCTTCCTGTGG-3’) were selected at exon 1 of *HaVipR2* according to the principle of 5’-N_20_NGG-3’ (underlined is the PAM sequence).

The forward and reverse overlapping oligonucleotides that contain the target DNA sequence (Target-F: 5’-TAATACGACTCACTATAGN_20_-3’; Target-R: 5’-TTCTAGCTCTAAAACN_20_-3’) were designed according to the manufacturer’s instructions (GeneArt™ Precision gRNA Synthesis Kit, Thermo Fisher Scientific, Shanghai, China).

Assembly PCR was used to generate the full-length gRNA DNA template. The PCR reaction mixture (25 µl) consisted of 12.5 µl of Phusion™ High-Fidelity PCR Master Mix (2x), 1 µl of Tracr Fragment + T7 Primer Mix, 1 µl of 0.3 μM Target F/R oligonucleotide mix, and 10.5 µl of Nuclease-free water. PCR was performed at 98 °C 10 s, 32 cycles of (98 °C 5 s, 55 °C 15 s), 72 °C 1 min and hold at 4 °C. The PCR products were then purified with FastPure Gel DNA Extraction Mini Kit (Vazyme, Nanjing, China). The sgRNA was synthesized by *in vitro* transcription utilizing the GeneArt™ Precision gRNA Synthesis Kit (Thermo Fisher Scientific) according to the manufacturer’s instructions.

#### Embryo microinjection

The collection and preparation of eggs were carried out as reported before (50). Briefly, fresh eggs laid within 2 hours were washed down from the gauzes in 0.5% sodium hypochlorite solution and rinsed with distilled water. After suction filtration, the eggs were lined up on a microscope slide (fixed with double-sided adhesive tape).

About one nanoliter mix of two sgRNAs (each 150 ng/μl) and Cas9 protein (300 ng/μl, TrueCut™ Cas9 Protein v2, Thermo Fisher Scientific) were injected into individual eggs using a FemtoJet and InjectMan NI 2 microinjection system (Eppendorf, Hamburg, Germany). The microinjection was completed within 2 h. Injected eggs were placed at 26 ± 1°C, 60 ± 10% RH for hatching.

#### Identification of HaVipR2 mutations mediated by CRISPR/Cas9

To identify the indel mutations at exon 1 of *HaVipR2*, a pair of diagnostic primers (F1: 5’-ATGACCAGAAAATATCAGTTGTGGG-3’, R1: 5-’GGCAAATTGGCTTCTCAGGACTAA-3’) was designed to amplify a 602-bp fragment flanking exon 1. To check for inherited mutations created by Cas9, genomic DNA from a pool of 10 second-instar larvae from the G_1_ progeny of each of the 14 single-pair families were prepared and directly sequenced. A cluster of double peaks around the Cas9 induced DSB cutting site indicated inherited mutations occurred in a single pair family. PCR products from family #13 were TA-cloned and sequenced to determine the exact indel mutation type. To establish a homozygous knockout strain, the G_1_ and G_2_ moths of family #13 were genotyped non-destructively before mating by removing one hind leg for PCR amplification and electrophoresis analysis of PCR products.

### Phylogenetic analysis of *HaVipR1* and *HaVipR2* in moths

Protein sequences of *H. armigera* HaVipR1 and HaVipR2 were used as queries in BLASTp searches against the full NCBI protein database. Subsequent hits were filtered to retain only sequences from Lepidoptera (Taxonomic ID: 7088). A single Dipteran sequence (*Anopheles stephensi*) for both HaVipR1 and HaVipR2 was included as an outgroup.

Protein sequences were aligned using MAFFT (v7.526; (51)) with default parameters. Maximum-likelihood phylogenies were inferred using IQ-TREE 2 via the online server (52), employing ModelFinder to select the best-fit amino-acid substitution model and ultrafast bootstrap support with 1,000 replicates. Trees were visualized and formatted in R using the phytools (v2.5-2; (53)) package, including cophylogenetic comparisons between HaVipR1 and HaVipR2 orthologs.

### Detection and characterisation of the transposable-element insertion within HaVipR2

#### Short-read SNP and INDEL analysis for the transposable element insertion in HaVipR2

The short-read alignments generated above for the F_0_, F_1_ and F_2_ individuals were evaluated with three variant callers: GATK (v4.2.0; (54)), FreeBayes, and bcftools mpileup/call.

For the GATK workflow, HaplotypeCaller was first run in per-sample mode to generate individual GVCFs, which were then combined and jointly genotyped using GenotypeGVCFs. For FreeBayes and BCFTools the tools were run with default parameters for all individuals. The insertion area on exon 5 was inspected visually with IGV to identify whether the caller produced the correct variant call and genotype, which is known based on the inheritance pattern.

FreeBayes and BCFTools did not produce a variant call at the insertion site. GATK was able to identify a small insertion (63bp) at the site for the F0 homozygous resistant and F1 heterozygous resistant individuals, but only ever genotypes this variant as heterozygous, even in both of the F_0_ individuals, where this is known to be homozygous.

#### Identification of the insertion using long-read assemblies

To identify structural variation within the resistance-associated interval, the *de novo* long-read assembly of the MIC F_0_ resistant individual, and an additional resistant individual from a different cross, was aligned to the GR reference genome using Minimap2 (v2.24). Alignments were sorted and indexed with samtools and manual inspection of the alignment in IGV revealed a large insertion within exon 5. The contig containing the insertion was extracted and annotated using RepeatMasker (Dfam 3.2 database) and the Repeat Finder tool in Geneious Prime (2023.2.1). Flanking sequences were examined to identify structural features, including inverted repeats and target-site duplication.

#### Assessment of transcription across the insertion

To evaluate transcriptional effects, RNA-seq reads from resistant individuals were mapped to the resistant assembly using Hisat2 (v2.2.1) with default parameters. The gene model for HaVipR2 was transferred to the resistant assembly using Geneious Prime. Alignments were visualized in IGV and splice junctions were assessed using Sashimi plots generated from the integrated RNA-seq tracks.

#### Evaluation of insertion detectability using short-read workflows

Short-read data from resistant individuals were aligned to the GR reference using BWA-MEM2 (v2.2.1). Processed BAM files (paired-end fixed, duplicates removed using Picard MarkDuplicates) were analysed for structural variants using Smoove (Pedersen 2020), which integrates LUMPY and SVTyper. Output VCFs were inspected in IGV for evidence of an insertion at HaVipR2.

Additional short-read insertion-detection tools were tested on the same alignments, including MindTheGap (v2.1), SCRAMBLE (v1.0), and Transposable Element Finder (TEF) (v1.1). TEF was run with the full repeat annotation generated by RepeatMasker. All results were manually inspected in IGV to see if the tools were able to identify the large insertion.

#### Evaluation using long-read alignment-based structural variant callers

Long-read datasets from resistant individuals were aligned to the GR reference using either Minimap2 (parameters: -x map-ont) or ngmlr (v0.2.7; default settings). Alignments were sorted and indexed with samtools, and three long-read SV callers were applied: SVIM (v1.4.2), NanoSV (v1.2.4), CuteSV (v1.0.12). Each program was run with default parameters, and resulting VCFs were assessed in IGV for structural variants at the HaVipR2 locus.

Mobile-element–oriented long-read callers TELR (v1.0) and TLDR (v1.1) were also run with default settings on the same alignments. Their output VCFs were similarly inspected for insertion calls.

#### Long-read assembly-based structural variant detection

As read-based tools did not identify the insertion, an assembly-based variant caller was tested. The *de novo* Flye assembly of the MIC F_0_ resistant individual was aligned to the GR reference using Minimap2 (-x asm20). These assembly-to-reference alignments were analysed using SVIM-asm (v1.0.0) with default parameters. Resulting variant predictions were visualised in IGV to confirm the presence and structure of the large insertion.

#### Assessment of insertion presence across Lepidoptera

The full insertion sequence was queried against the NCBI nr/nt database using BLASTn (default parameters) to identify homologous sequences in other Lepidoptera genomes. Hits were reviewed for TE family assignment and sequence similarity.

### Re-annotation of genomes lacking HaVipR2

For species in which HaVipR2 was absent from the published protein annotation (Papilio polytes (GCF_000836215.1) and *Ostrinia nubilalis* (GCF_963855985.1)), the corresponding genome assemblies and RefSeq annotations were downloaded from NCBI. The *H. armigera* HaVipR1 and HaVipR2 protein sequences were provided as queries to MetaEuk (default parameters), which was run directly on each downloaded genome assembly to identify homologous loci independent of the existing annotation. The MetaEuk annotations for HaVipR1 and HaVipR2 were then inspected manually in IGV to assess validity.

## Acknowledgments

This work was supported by the Commonwealth Scientific and Industrial Research Organisation (CSIRO) through the Environomics Future Science Platform (FSP) (PhD Top-Up support to A.B.) and the Advanced Engineering Biology FSP (Post-Doctoral support).

Funding was also provided by the Cotton Research and Development Corporation (CRDC) through projects CSE1801 and CSP2204.

## Author contributions

Conceptualization:

AB, TW, AP, KG, YW, TT

Methodology:

AB, ES, DA, YW, MW, BJ, CGF

Investigation:

AB, LW, CGF, MW, BJ

Visualization:

AB

Supervision:

DA, ES, TW, AP

Writing-original draft:

AB, TW, AP

Writing-review & editing:

AB, KG, YW

Senior authors:

TW, AP

## Data availability

Sequencing data are available under BioProject PRJNA1074112 (Long-Read+ShortReads) and BioProject PRJNA1119665 (short-reads of full MIC and FIC + RNA of 17-294 and GR). The reference genome (susceptible lab line) generated in this study is available on GenBank (GCA_040954515.1).

## References

1. Tay WT, Soria MF, Walsh T, Thomazoni D, Silvie P, Behere GT, et al. A Brave New World for an Old World Pest: Helicoverpa armigera (Lepidoptera: Noctuidae) in Brazil. Knapp M, editor. PLoS One. 2013 Nov 18;8(11):e80134. doi:10.1371/journal.pone.0080134

2. Zafar MM, Razzaq A, Farooq MA, Rehman A, Firdous H, Shakeel A, et al. Insect resistance management in Bacillus thuringiensis cotton by MGPS (multiple genes pyramiding and silencing). Journal of Cotton Research. 2020 Dec 20;3(1):33. doi:10.1186/s42397-020-00074-0

3. Lu Y, Wu K, Jiang Y, Guo Y, Desneux N. Widespread adoption of Bt cotton and insecticide decrease promotes biocontrol services. Nature. 2012 Jul 19;487(7407):362–5. doi:10.1038/nature11153

4. Wells L. New generation Bollgard III to blitz cotton plantings [Industry [Internet]. Grain Central. Available from: https://www.graincentral.com/cropping/new-generation-bollgard-iii-to-blitz-cotton-plantings/

5. Jouzani GS, Valijanian E, Sharafi R. Bacillus thuringiensis: a successful insecticide with new environmental features and tidings. Appl Microbiol Biotechnol. 2017 Apr 24;101(7):2691–711. doi:10.1007/s00253-017-8175-y

6. Jurat-Fuentes JL, Heckel DG, Ferré J. Mechanisms of Resistance to Insecticidal Proteins from Bacillus thuringiensis. Annu Rev Entomol. 2021 Jan 7;66(1):121–40. doi:10.1146/annurev-ento-052620-073348

7. Melo AL de A, Soccol VT, Soccol CR. *Bacillus thuringiensis* : mechanism of action, resistance, and new applications: a review. Crit Rev Biotechnol. 2016 Mar 3;36(2):317–26. doi:10.3109/07388551.2014.960793

8. Shan Y, Jin M, Chakrabarty S, Yang B, Li Q, Cheng Y, et al. Sf-FGFR and Sf-SR-C Are Not the Receptors for Vip3Aa to Exert Insecticidal Toxicity in Spodoptera frugiperda. Insects. 13(6). doi:10.3390/insects13060547

9. Jin M, Shan Y, Peng Y, Wang W, Zhang H, Liu K, et al. Downregulation of a transcription factor associated with resistance to Bt toxin Vip3Aa in the invasive fall armyworm. Proceedings of the National Academy of Sciences. 2023 Oct 31;120(44):2306932120. doi:10.1073/pnas.2306932120

10. Liu Z, Liao C, Zou L, Jin M, Shan Y, Quan Y, et al. Retrotransposon-mediated disruption of a chitin synthase gene confers insect resistance to Bacillus thuringiensis Vip3Aa toxin. PLoS Biol. 2024 Jul 1;22(7):e3002704. doi:10.1371/JOURNAL.PBIO.3002704 PubMed PMID: 38954724.

11. Wang P, Liu Z, Kang Q, Liao C, Zou L, Mao K, et al. Functional loss of CHS2 confers high-level resistance to Bacillus thuringiensis Vip3Aa in Spodoptera exigua and Agrotis ipsilon. Pest Manag Sci. 2025. doi:10.1002/PS.70226

12. Heu CC, Schutze IX, LeRoy DM, Wang YH, DeGain BA, Kerns DD, et al. Knockout of chitin synthase gene confers resistance to Bt toxin Vip3Aa in Helicoverpa zea. Pest Manag Sci. 2025. doi:10.1002/PS.70248

13. Bachler A, Padovan A, Anderson CJ, Wei Y, Wu Y, Pearce S, et al. Disruption of HaVipR1 confers Vip3Aa resistance in the moth crop pest Helicoverpa armigera. PLoS Biol. 2025 May 1;23(5):e3003165. doi:10.1371/JOURNAL.PBIO.3003165 PubMed PMID: 40440223.

14. Zhang Z, Pang X, Wang L, Tay WT, Gordon KHJ, Walsh TK, et al. Knockout of the SfVipR1 gene confers high-level resistance to Bacillus thuringiensis Vip3Aa toxin in Spodoptera frugiperda. Pest Manag Sci. 2025. doi:10.1002/PS.70190 PubMed PMID: 40899875.

15. Pezzini D, Taylor KL, Reisig DD, Fritz ML. Cross-pollination in seed-blended refuge and selection for Vip3A resistance in a lepidopteran pest as detected by genomic monitoring. Proceedings of the National Academy of Sciences. 2024 Mar 26;121(13):e2319838121. doi:10.1073/PNAS.2319838121 PubMed PMID: 38513093.

16. Yang F, Head GP, Kerns DD, Jurat-Fuentes JL, Santiago-González JC, Kerns DL. Diverse genetic basis of Vip3Aa resistance in five independent field-derived strains of Helicoverpa zea in the US. Pest Manag Sci. 2024 Jun 1;80(6):2796–803. doi:10.1002/PS.7988 PubMed PMID: 38327120.

17. Gahan LJ, Gould F, Heckel DG. Identification of a Gene Associated with Bt Resistance in *Heliothis virescens*. Science (1979). 2001 Aug 3;293(5531):857–60. doi:10.1126/science.1060949

18. Tossolini I, Mencia R, Arce AL, Manavella PA. The genome awakens: transposon-mediated gene regulation. Trends Plant Sci. 2025 Aug 1;30(8):857–71. doi:10.1016/J.TPLANTS.2025.02.005 PubMed PMID: 40069082.

19. Mahmoud M, Gobet N, Cruz-Dávalos DI, Mounier N, Dessimoz C, Sedlazeck FJ. Structural variant calling: The long and the short of it. Genome Biol. 2019 Nov 20;20(1):1–14. doi:10.1186/S13059-019-1828-7/TABLES/2 PubMed PMID: 31747936.

20. Fabrick JA, Wu Y. Mechanisms and molecular genetics of insect resistance to insecticidal proteins from Bacillus thuringiensis. Adv In Insect Phys. 2023 Jan 1;65:123–83. doi:10.1016/BS.AIIP.2023.09.005

21. Heller D, Vingron M. SVIM-asm: structural variant detection from haploid and diploid genome assemblies. Robinson P, editor. Bioinformatics. 2021 Apr 1;36(22–23):5519–21. doi:10.1093/bioinformatics/btaa1034

22. Daron J, Bergman A, Lambrechts L. Dynamics and evolution of transposable elements in mosquito genomes. Curr Opin Insect Sci. 2025 Oct 1;71:101406. doi:10.1016/J.COIS.2025.101406

23. Novinec M, Kordiš D, Turk V, Lenarčič B. Diversity and Evolution of the Thyroglobulin Type-1 Domain Superfamily. Mol Biol Evol. 2006 Apr 1;23(4):744–55. doi:10.1093/molbev/msj082

24. Mihelič M, Turk D. Two decades of thyroglobulin type-1 domain research. Biol Chem. 2007 Nov 1;388(11):1123–30. doi:10.1515/BC.2007.155 PubMed PMID: 17976004.

25. Castro-Gomes T, Corrotte M, Tam C, Andrews NW. Plasma Membrane Repair Is Regulated Extracellularly by Proteases Released from Lysosomes. Gasman S, editor. PLoS One. 2016 Mar 30;11(3):e0152583. doi:10.1371/journal.pone.0152583

26. Hou X, Han L, An B, Cai J. Autophagy induced by Vip3Aa has a pro-survival role in *Spodoptera frugiperda* Sf9 cells. Virulence. 2021 Dec 31;12(1):509–19. doi:10.1080/21505594.2021.1878747

27. Hou X, Han L, An B, Zhang Y, Cao Z, Zhan Y, et al. Mitochondria and Lysosomes Participate in Vip3Aa-Induced Spodoptera frugiperda Sf9 Cell Apoptosis. Toxins (Basel). 2020;12(2):116. doi:10.3390/TOXINS12020116 PubMed PMID: 32069858.

28. Gomis-Cebolla J, Wang Y, Quan Y, He K, Walsh T, James B, et al. Analysis of cross-resistance to Vip3 proteins in eight insect colonies, from four insect species, selected for resistance to Bacillus thuringiensis insecticidal proteins. J Invertebr Pathol. 2018 Jun;155:64–70. doi:10.1016/j.jip.2018.05.004

29. Apirajkamol N, James B, Gordon KHJ, Walsh TK, McGaughran A. Oxidative stress delays development and alters gene expression in the agricultural pest moth, Helicoverpa armigera. Ecol Evol. 2020 Jun 1;10(12):5680–93. doi:10.1002/ECE3.6308;PAGE:STRING:ARTICLE/CHAPTER

30. Heckel DG, Gahan LJ, Liu YB, Tabashnik BE. Genetic mapping of resistance to Bacillus thuringiensis toxins in diamondback moth using biphasic linkage analysis. Proceedings of the National Academy of Sciences. 1999 Jul 20;96(15):8373–7. doi:10.1073/pnas.96.15.8373

31. Yang F, González JCS, Little N, Reisig D, Payne G, Dos Santos RF, et al. First documentation of major Vip3Aa resistance alleles in field populations of Helicoverpa zea (Boddie) (Lepidoptera: Noctuidae) in Texas, USA. Scientific Reports 2020 10:1. 2020 Apr 3;10(1):5867–. doi:10.1038/s41598-020-62748-8 PubMed PMID: 32246037.

32. Yang F, Head GP, Price PA, Santiago González JC, Kerns DL. Inheritance of <scp> *Bacillus thuringiensis* Cry2Ab2 </scp> protein resistance in <scp> *Helicoverpa zea* </scp> (Lepidoptera: Noctuidae). Pest Manag Sci. 2020 Nov 8;76(11):3676–84. doi:10.1002/ps.5916

33. Koren S, Walenz BP, Berlin K, Miller JR, Bergman NH, Phillippy AM. Canu: scalable and accurate long-read assembly via adaptive *k*-mer weighting and repeat separation. Genome Res. 2017 May;27(5):722–36. doi:10.1101/gr.215087.116

34. Walker BJ, Abeel T, Shea T, Priest M, Abouelliel A, Sakthikumar S, et al. Pilon: An Integrated Tool for Comprehensive Microbial Variant Detection and Genome Assembly Improvement. Wang J, editor. PLoS One. 2014 Nov 19;9(11):e112963. doi:10.1371/journal.pone.0112963

35. Roach MJ, Schmidt SA, Borneman AR. Purge Haplotigs: allelic contig reassignment for third-gen diploid genome assemblies. BMC Bioinformatics 2018 19:1. 2018 Nov 29;19(1):460–. doi:10.1186/S12859-018-2485-7 PubMed PMID: 30497373.

36. Zhou C, McCarthy SA, Durbin R. YaHS: yet another Hi-C scaffolding tool. Bioinformatics. 2023 Jan 1;39(1). doi:10.1093/BIOINFORMATICS/BTAC808 PubMed PMID: 36525368.

37. Jain C, Koren S, Dilthey A, Phillippy AM, Aluru S. A fast adaptive algorithm for computing whole-genome homology maps. Bioinformatics. 2018 Sep 1;34(17):i748–56. doi:10.1093/BIOINFORMATICS/BTY597 PubMed PMID: 30423094.

38. Allio R, Schomaker-Bastos A, Romiguier J, Prosdocimi F, Nabholz B, Delsuc F. MitoFinder: Efficient automated large-scale extraction of mitogenomic data in target enrichment phylogenomics. Mol Ecol Resour. 2020 Jul 1;20(4):892. doi:10.1111/1755-0998.13160 PubMed PMID: 32243090.

39. Flynn JM, Hubley R, Goubert C, Rosen J, Clark AG, Feschotte C, et al. RepeatModeler2 for automated genomic discovery of transposable element families. Proc Natl Acad Sci U S A. 2020 Apr 28;117(17):9451–7. doi:10.1073/PNAS.1921046117;PAGE:STRING:ARTICLE/CHAPTER PubMed PMID: 32300014.

40. Shumate A, Salzberg SL. Liftoff: accurate mapping of gene annotations. Bioinformatics. 2021 Jul 19;37(12):1639–43. doi:10.1093/BIOINFORMATICS/BTAA1016 PubMed PMID: 33320174.

41. Jones P, Binns D, Chang HY, Fraser M, Li W, McAnulla C, et al. InterProScan 5: genome-scale protein function classification. Bioinformatics. 2014 May 1;30(9):1236. doi:10.1093/BIOINFORMATICS/BTU031 PubMed PMID: 24451626.

42. Kolmogorov M, Yuan J, Lin Y, Pevzner PA. Assembly of long, error-prone reads using repeat graphs. Nat Biotechnol. 2019 May 1;37(5):540–6. doi:10.1038/s41587-019-0072-8

43. Md V, Misra S, Li H, Aluru S. Efficient architecture-aware acceleration of BWA-MEM for multicore systems. Proceedings - 2019 IEEE 33rd International Parallel and Distributed Processing Symposium, IPDPS 2019. 2019 May 1;314–24. doi:10.1109/IPDPS.2019.00041

44. Danecek P, Bonfield JK, Liddle J, Marshall J, Ohan V, Pollard MO, et al. Twelve years of SAMtools and BCFtools. Gigascience. 2021 Jan 29;10(2). doi:10.1093/gigascience/giab008

45. Garrison E, Marth G. Haplotype-based variant detection from short-read sequencing.

46. Kim D, Paggi JM, Park C, Bennett C, Salzberg SL. Graph-based genome alignment and genotyping with HISAT2 and HISAT-genotype. Nat Biotechnol. 2019 Aug 2;37(8):907–15. doi:10.1038/s41587-019-0201-4

47. Pertea M, Pertea GM, Antonescu CM, Chang TC, Mendell JT, Salzberg SL. StringTie enables improved reconstruction of a transcriptome from RNA-seq reads. Nat Biotechnol. 2015 Mar 18;33(3):290–5. doi:10.1038/nbt.3122

48. Love MI, Huber W, Anders S. Moderated estimation of fold change and dispersion for RNA-seq data with DESeq2. Genome Biology 2014 15:12. 2014 Dec 5;15(12):550–. doi:10.1186/S13059-014-0550-8 PubMed PMID: 25516281.

49. Yang YH, Yang YJ, Gao WY, Guo JJ, Wu YH, Wu YD. Introgression of a disrupted cadherin gene enables susceptible Helicoverpa armigera to obtain resistance to Bacillus thuringiensis toxin Cry1Ac. Bull Entomol Res. 2009 Apr;99(2):175–81. doi:10.1017/S0007485308006226 PubMed PMID: 18954492.

50. Wang J, Zhang H, Wang H, Zhao S, Zuo Y, Yang Y, et al. Functional validation of cadherin as a receptor of Bt toxin Cry1Ac in Helicoverpa armigera utilizing the CRISPR/Cas9 system. Insect Biochem Mol Biol. 2016 Sep 1;76:11–7. doi:10.1016/j.ibmb.2016.06.008 PubMed PMID: 27343383.

51. Katoh K, Standley DM. MAFFT Multiple Sequence Alignment Software Version 7: Improvements in Performance and Usability. Mol Biol Evol. 2013 Apr;30(4):772. doi:10.1093/MOLBEV/MST010 PubMed PMID: 23329690.

52. Trifinopoulos J, Nguyen LT, von Haeseler A, Minh BQ. W-IQ-TREE: a fast online phylogenetic tool for maximum likelihood analysis. Nucleic Acids Res. 2016 Jul 8;44(W1):W232–5. doi:10.1093/NAR/GKW256 PubMed PMID: 27084950.

53. Revell LJ. phytools 2.0: an updated R ecosystem for phylogenetic comparative methods (and other things). PeerJ. 2024;12:e16505. doi:10.7717/PEERJ.16505 PubMed PMID: 38192598.

54. McKenna A, Hanna M, Banks E, Sivachenko A, Cibulskis K, Kernytsky A, et al. The Genome Analysis Toolkit: A MapReduce framework for analyzing next-generation DNA sequencing data. Genome Res. 2010 Sep;20(9):1297. doi:10.1101/GR.107524.110 PubMed PMID: 20644199.

