## Supplementary Information for "Transposable element disruption of a second thyroglobulin-like gene confers Vip3Aa resistance in *Helicoverpa armigera*"

OFFICIAL

Science Advances
AAAS

Supplementary Materials for

### Transposable element disruption of a second thyroglobulin-like gene confers Vip3Aa resistance in *Helicoverpa armigera*

Andreas Bachler *et al.*

**This PDF file includes:**

Supplementary Text
Figs. S1 to S6
Tables S1 to S7



OFFICIAL

**Supplementary Section 1**

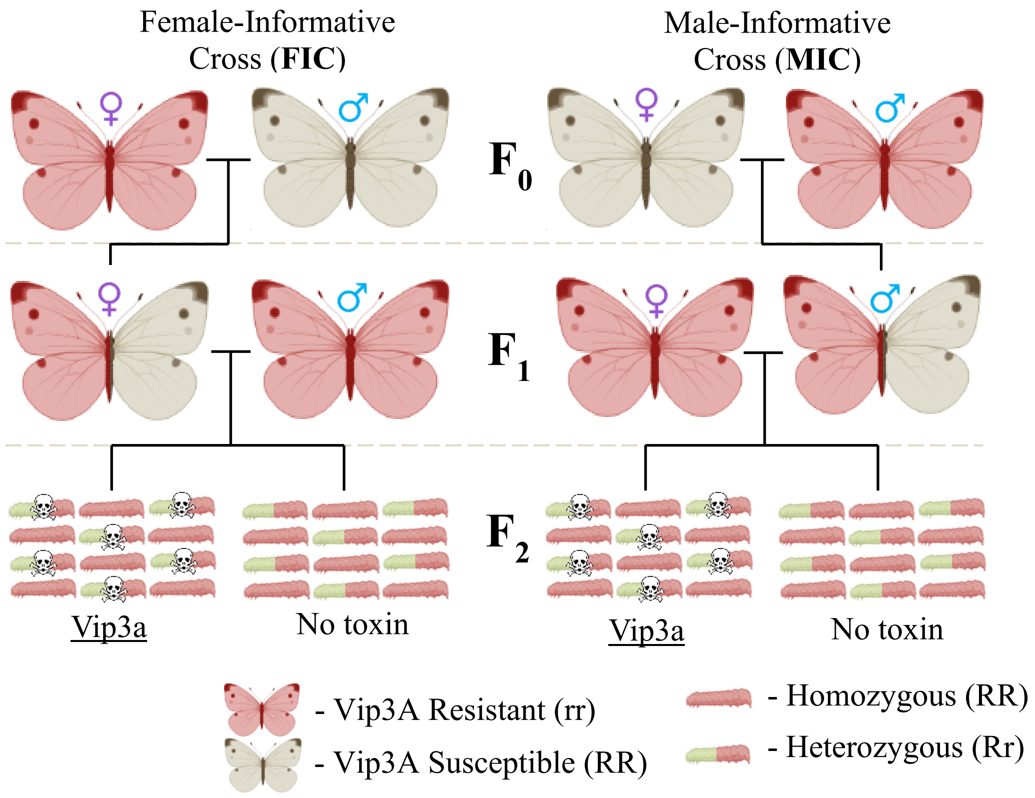


**Figure S1. Schematic of resistance crossing method for assessing linkage of Vip3A resistance in the resistant line 17-294.**
Multiple single pair matings were conducted with both sexes as the resistant individual to allow identification of the chromosome associating with resistance (through Female-Informative Crosses, or FIC) and the specific region of a chromosome (through Male-Informative Crosses, or MIC). F2 offspring were either exposed to Vip3A surface treatment or not (control). All samples in an MIC and FIC cross were sequenced using short-read sequencing and the 4 MIC $F_0$ and $F_1$ individuals were additionally processed for long-read sequencing (see **Supplementary Table S1**).

**Table S1. Overview of samples chosen for sequencing from sex-based crosses evaluated.** Long and short-read data was uploaded to SRA at PRJNA1074112.

| Cross | Sample | Short-read sequenced | Long-read sequenced |
|---|---|---|---|
| **Male-Informative Cross (MIC)** | F0 Male and Female | ✓ | ✓ |
| | F1 Male and Female | ✓ | ✓ |
| | F2 water-treated offspring (24x) | ✓ | ✗ |
| | F2 Vip3A-treated offspring (24x) | ✓ | ✗ |
| **Female-Informative Cross (FIC)** | F0 Male and Female | ✓ | ✗ |
| | F1 Male and Female | ✓ | ✗ |
| | F2 water-treated offspring (24x) | ✓ | ✗ |
| | F2 Vip3A-treated offspring (24x) | ✓ | ✗ |
| **Follow-up Male-Informative Cross** | F0 Male (male from 17-294 resistant line) "Follow-up" sample | ✓ | ✓ |

**Table S2. Differential expression of *Helicoverpa armigera* orthologs of candidate genes previously implicated in Vip3A resistance in *Spodoptera frugiperda*.** Expression differences between the resistant ("*X17*") and susceptible (*GR*) strains are shown for genes identified by Jin et al. (2023). Values include mean raw counts for each strain, $log_2$ fold change, and adjusted *p*-values. Genes with significant differential expression (*adjusted* $p < 0.01$) are highlighted in bold.

| ***HELICOVERPA ARMIGERA*** **GENEID** | **GENE NAME** | **GENE NAME SHORT** | **MEAN (RAW COUNTS) SUSCEPTIBLE / "GR"** | **MEAN (RAW COUNTS) RESIST / "X17"** | **LOG2 FOLD CHANGE** | **ADJUSTED P-VALUE** |
|---|---|---|---|---|---|---|
| LOC110370773 | phenoloxidase 1 | "P02" | 4336 | 5174 | -0.46 | 0.663 |
| LOC110371189 | phenoloxidase-activating enzyme | PAE | 909 | 1238 | -0.96 | 0.156 |
| LOC110372413 | LDL-receptor class A domain-containing protein 1 / "Scavenger receptor-C" | SRC | 581 | 611 | -0.67 | 0.32 |
| **LOC110373728** | **fibroblast growth factor receptor homolog 1** | **FGFR** | **343** | **7509** | **4.34** | **<0.00000001** |
| **LOC110374447** | **autophagy protein 5** | **ATG5** | **625** | **2470** | **1.89** | **0.006** |
| LOC110377400 | 40S ribosomal protein S2 | S2 | 11844 | 9143 | -1.63 | 0.046 |
| LOC126053489 | myb protein-like | SfMyb | 90 | 332 | 1.22 | 0.054 |
| LOC110371908 | chitin synthase-2 | SfCHS2 | 10429 | 15255 | -0.45 | 0.565 |

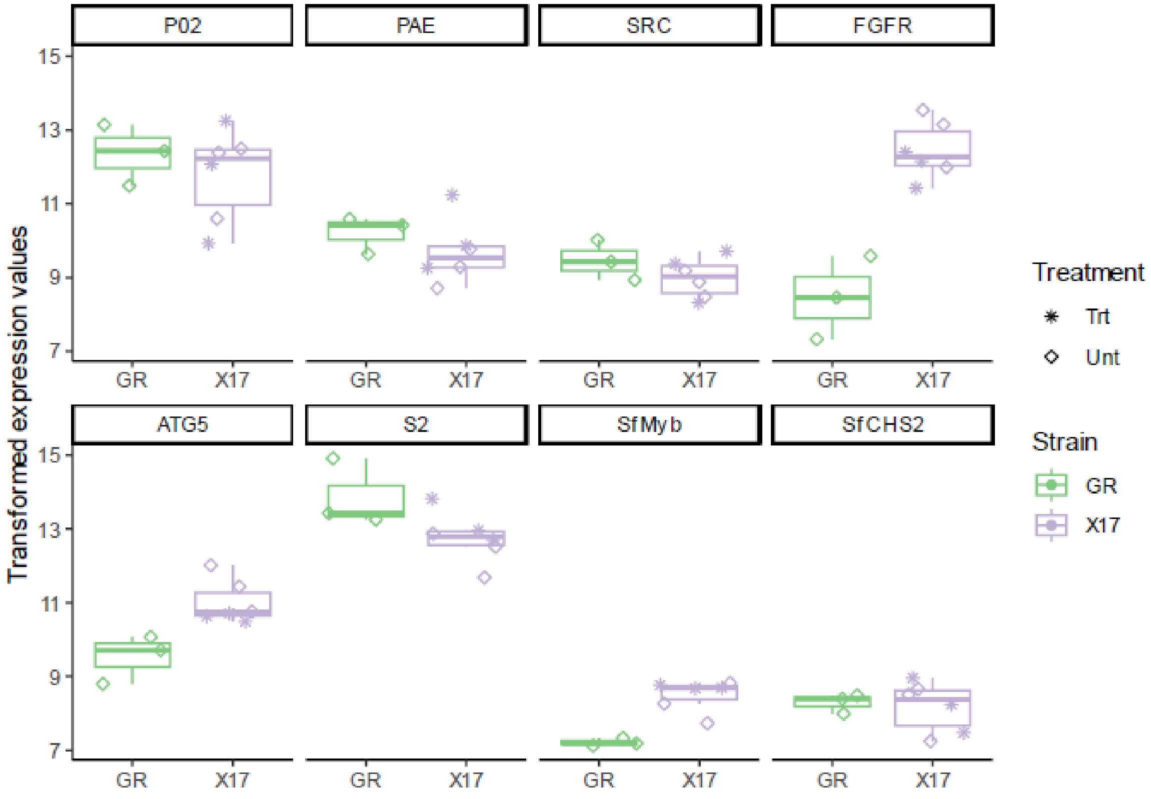


**Figure S2. Expression of candidate genes implicated in Vip3A resistance.** Variance-stabilized count values (vst, DESeq2) are shown for genes previously reported to interact with or modulate Vip3A toxicity in *Spodoptera frugiperda*. Data are presented for all biological replicates of the resistant (*X17*) and susceptible (*GR*) *H. armigera* lines, with outlier points removed from the boxplots (all individual values are plotted). None of these genes were identified in the resistance-associated region. Raw differential expression statistics are provided in Supplementary Table S2.

**Table S3. List of the top 10 most differentially expressed genes in the Vip3A resistant line in the target region associating with resistance.**

| GeneID | GeneName | Regulation | Presumed function |
|---|---|---|---|
| **LOC126056805** | thyroglobulin-like protein | Down | Protease regulation |
| **LOC110370202** | epimerase family protein SDR39U1 | Down | Metabolic pathway |
| **LOC110370137** | ganglioside-induced differentiation-associated protein 1 | Down | Cell differentiation |
| **LOC110370098** | cytochrome b-c1 complex subunit 2, mitochondrial | Down | Cellular respiration/ mitochondrial electron transport chain |
| **LOC110379619** | peroxisomal membrane protein PMP34 | Down | Lipid transport |
| **LOC110370091** | serine/threonine-protein kinase RIO1 | Up | Protein phosphorylation, cellular regulation |
| **LOC110370196** | programmed cell death protein 5 | Up | Cell regulation and apoptosis |
| **LOC110370072** | purine nucleoside phosphorylase | Up | Nucleotide synthesis |
| **LOC110370044** | enoyl-CoA delta isomerase 2 | Up | Lipid metabolism |
| **LOC110370043** | uncharacterized protein | Up | BLAST similarity to TESPA1, cell regulation and signalling |

**Table S4 List of all significantly differentially expressed genes (adjusted p-value <0.01) arising from exposure to Vip3A treatment of the resistant line.**
Output from DeSeq2 analysis of RNA-Seq alignments using Hisat2 and counts derived from Stringtie. Up and down regulation indicated in blue and red.

| Ref | baseMean | log2FoldChange | Regulation | lfcSE | padj | HarmChrom | trans_id | ref_gene_id | GeneName | Function |
|---|---|---|---|---|---|---|---|---|---|---|
| **LOC110371128** | 30617.61001 | 4.058944986 | Up | 0.839412 | 1.99E-06 | scaffold_28_RagTag | rna-XM_021327242.2 | gene-LOC110371128 | uncharacterized LOC110371128 | Similarity to midasin and potential target of myb but not from PNAS paper |
| **LOC110375614** | 6917.998895 | -2.621266503 | Down | 0.517321 | 7.16E-07 | scaffold_16_RagTag | rna-XM_021333791.2 | gene-LOC110375614 | phospholipase A2 | |
| **LOC110382852** | 4351.530161 | 3.782355297 | Up | 0.623799 | 3.71E-09 | scaffold_12_RagTag | rna-XM_021343556.2 | gene-LOC110382852 | protein croquemort | |
| **LOC110372893** | 3807.907787 | -6.678779899 | Down | 0.461924 | 8.63E-46 | scaffold_05_RagTag | rna-XM_049836228.1 | gene-LOC110372893 | kinesin-like protein KIF13A | |
| **LOC110383098** | 1495.925378 | -3.801733604 | Down | 0.499457 | 1.32E-13 | scaffold_15_RagTag | rna-XM_049842790.1 | gene-LOC110383098 | YLP motif-containing protein 1 | |
| **LOC110376466** | 1216.558896 | 2.46609312 | Up | 0.469275 | 3.04E-07 | scaffold_18_RagTag | rna-XM_021334951.2 | gene-LOC110376466 | uncharacterized LOC110376466 | |
| **LOC110377108** | 845.6599301 | 1.621483481 | Up | 0.340017 | 2.41E-06 | scaffold_20_RagTag | rna-XM_021335806.2 | gene-LOC110377108 | zinc finger CCCH domain-containing protein 13 | |
| **LOC126055067** | 781.6211786 | 4.627980258 | Up | 0.661956 | 1.18E-11 | scaffold_15_RagTag | rna-XM_049842668.1 | gene-LOC126055067 | surfeit locus protein 4 homolog | |
| **LOC110384138** | 714.9000126 | -5.521454488 | Down | 1.079297 | 6.09E-07 | scaffold_22_RagTag | rna-XM_021345249.2 | gene-LOC110384138 | eukaryotic translation initiation factor 3 subunit G | |
| **LOC110376609** | 647.825113 | 4.418879044 | Up | 0.359534 | 1.99E-33 | scaffold_20_RagTag | rna-XM_021335157.2 | gene-LOC110376609 | caspase-8 | Apoptosis and cell regulation |
| **LOC110373074** | 643.196717 | 5.018682322 | Up | 0.932529 | 1.60E-07 | scaffold_23_RagTag | rna-XM_049847495.1 | gene-LOC110373074 | pre-mRNA-splicing factor Slu7 | |
| **LOC110378224** | 486.3987156 | -2.435397151 | Down | 0.564116 | 1.67E-05 | scaffold_01_RagTag | rna-XM_049841218.1 | gene-LOC110378224 | protein ABHD13 | |
| **LOC110383188** | 386.9435059 | -2.103824068 | Down | 0.440418 | 2.39E-06 | scaffold_08_RagTag | rna-XM_049838072.1 | gene-LOC110383188 | SPRY domain-containing SOCS box protein 3 | |
| **LOC110369913** | 270.540069 | 2.297514888 | Up | 0.480838 | 2.39E-06 | scaffold_10_RagTag | rna-XM_021325547.2 | gene-LOC110369913 | sodium-coupled monocarboxylate transporter 1 | |
| **LOC110376578** | 224.7887454 | 6.050861677 | Up | 1.279893 | 2.77E-06 | scaffold_15_RagTag | rna-XM_049841747.1 | gene-LOC110376578 | probable phospholipid-transporting ATPase IIB | |
| **LOC110371920** | 216.4236228 | 2.153161017 | Up | 0.438753 | 1.44E-06 | scaffold_04_RagTag | rna-XM_049835769.1 | gene-LOC110371920 | CD109 antigen | |
| **LOC110374584** | 195.5261902 | 2.268306177 | Up | 0.533125 | 2.15E-05 | scaffold_01_RagTag | rna-XM_021332347.2 | gene-LOC110374584 | proton-coupled amino acid transporter-like protein CG1139 | |
| **LOC110372522** | 194.6262388 | -3.406064792 | Down | 0.785324 | 1.56E-05 | scaffold_23_RagTag | rna-XM_021329288.2 | gene-LOC110372522 | uncharacterized LOC110372522 | |
| **LOC110371421** | 163.8572084 | 10.66225337 | Up | 1.36867 | 5.22E-14 | scaffold_31_RagTag | rna-XM_049839173.1 | gene-LOC110371421 | uncharacterized LOC110371421 | |
| **LOC110377873** | 133.2754904 | 10.36394541 | Up | 1.57171 | 1.39E-10 | scaffold_23_RagTag | rna-XM_049847573.1 | gene-LOC110377873 | formin-binding protein 4 | |
| **LOC110374446** | 121.8284757 | -5.515575389 | Down | 0.716003 | 8.62E-14 | scaffold_15_RagTag | rna-XM_049842526.1 | gene-LOC110374446 | torsin-1A | |
| **LOC110370899** | 99.67472348 | -4.553181658 | Down | 0.892886 | 6.33E-07 | scaffold_15_RagTag | rna-XM_021326906.2 | gene-LOC110370899 | zinc finger matrin-type protein 3 | |
| **LOC110384623** | 54.64396655 | -5.880746236 | Down | 1.225316 | 2.30E-06 | scaffold_02_RagTag | rna-XM_049842884.1 | gene-LOC110384623 | dymeclin | |
| **LOC126054795** | 41.63692107 | 8.686195851 | Up | 1.593777 | 1.23E-07 | scaffold_02_RagTag | rna-XM_049841537.1 | gene-LOC126054795 | chorion class A protein Ld2/Ld41-like | |

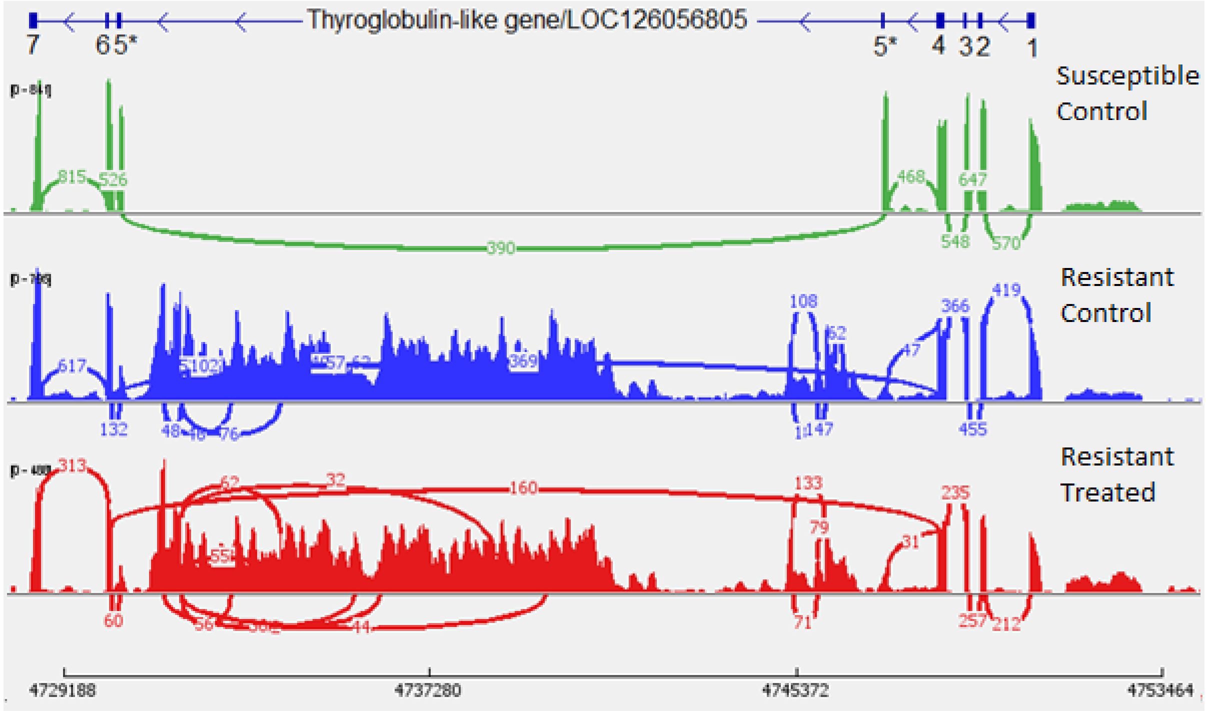


**Figure S5. Sashimi plot of RNA-Seq mapping indicates disruption of exon 5 of the thyroglobulin-like gene LOC126056805 in the Vip3A resistant line.** RNA sequence data was mapped to the resistant reference using Hisat2 and visualised in IGV. Gene annotations of the thyroglobulin-like gene were generated using metaeuk, with exon 5 indicated as split by the large insertion, and is provided with the exon numbers indicated. Splice junctions with minimum read support of 30 are shown. The "Susceptible Control" individual (first track, green) is from the "GR" line with $H_2O$ treatment, the "Resistant Control" individual (second track, blue) is from the Vip3A resistant line with $H_2O$ treatment and the "Resistant Treated" individual (third track, red) is from the Vip3A resistant line with Vip3A treatment.

**Supplementary Section 2**
**Transposable element insertion assessment – Methods**

Short-read variant identification

Short-read genomic data from all individuals in the Male-Informative Cross (MIC) and Female-Informative Cross (FIC) were aligned to the *H. armigera* reference genome (GCA_040954515.1; generated as part of this study) using BWA-MEM2 (v2.2.1; Md et al., 2019) with default parameters. Pairing information was corrected using samtools fixmate (v1.18; Danecek et al., 2021), duplicates were removed using Picard MarkDuplicates, and alignments were then sorted and indexed with samtools.

Variant calling was performed using GATK HaplotypeCaller (v4.2.0) for all $F_0$ and $F_1$ individuals, followed by joint genotyping across all samples. Variants were subsequently filtered to retain only those consistent with the expected monogenic recessive inheritance pattern - i.e., homozygous-alternate in both '17-294' resistant $F_0$ individuals, homozygous-reference in the susceptible parent, and heterozygous in the $F_1$ hybrids.

Functional effects of retained variants were annotated using SnpEff (v5.0e; Cingolani et al., 2012). All putative disruptive variants were then cross-checked against two known susceptible *H. armigera* assemblies (Harm1.0/GCA_002156985.1 and HaSCD2/ GCA_023701775.1) by aligning those assemblies to the study reference genome and inspecting the corresponding nucleotide positions.

Short-read structural variant detection

Structural variant analysis was carried out using Smoove which carries out optimization of Lumpy and a range of other tools to identify and genotype a range of structural variants. Genotyping accuracy of structural variant callers is not as reliable as variant genotyping and so a looser filter was applied, requiring at least 1 non-susceptible individual to have a SV intersect with a coding-sequence (CDS), 5' Untranslated Region (5' UTR) or 3' Untranslated Region (3' UTR). Candidate SVs were annotated using gene models from the reference genome and compared across resistant and susceptible individuals for consistency with recessive inheritance. SVs were also validated against the Harm1.0 and HaSCD2 susceptible assemblies as previously.

Long-read structural variant detection

Long-read (ONT/PacBio) data from all MIC individuals were aligned to the reference genome using Minimap2 (v2.24). Structural variants were identified using Sniffles (v.2.7.0; Sedlazeck et al., 2018)).

#### Transposable element insertion assessment - Results

Short-read variant identification

Across the 3.5 Mb resistance-linked region (250 genes), nine SnpEff-annotated "HIGH" impact variants were detected. However, inspection against susceptible assemblies showed that most were also present in susceptible genomes. Several apparent stop-gained SNPs were actually mis-annotated due to adjacent SNP/indel context producing missense rather than nonsense changes. Remaining frameshifts were resolved by paired compensatory indels or lacked consistent read support. No genuine coding disruptions matching the recessive inheritance pattern were identified using short-read variant calling.

**Table S5. Annotated variants in resistance linked region with putative disruption as identified by SnpEff.**

| GeneID | GeneProduct | Variant type | Disruption type | Comment. *Linked to resistance?* |
|---|---|---|---|---|
| LOC110370052 | Mismatch repair endonuclease PMS2 | 2bp Del | Frameshift | Highly variable region in HARM1.0 reference, large deletion identified. Highly variable in HaSCD reference, many small insertions and deletions identified. *Not linked to resistance.* |
| | | 2bp Ins | Frameshift | Matching deletion identified in HaSCD reference. *Not linked to resistance.* |
| | | SNP | Stop gained | Adjacent SNP present, correct call is an MNP not a SNP. These variants should be considered together, causes a mis-sense mutation (S154Y) not a stop-gained event. *Not linked to resistance.* |
| LOC110370193 | Uncharacterized LOC110370193 | 16bp Del | Splice site acceptor | Matching deletion identified in HaSCD reference. *Not linked to resistance.* |
| LOC110370191 | Uncharacterized LOC110370191 | 4bp Ins | Frameshift | Matching insertion seen in HaSCD and HARM1.0 reference. *Not linked to resistance.* |
| LOC110370182 | Dynein intermediate chain 3, ciliary-like | SNP | Splice site donor | HaSCD and HARM1.0 reference support SNP, potential alternative splice site. *Not linked to resistance.* |
| LOC110370211 | Calcium-binding protein P-like | 2bp Del | Frameshift | With below adjacent small INDEL reading frame restored. *Not linked to resistance.* |
| | | 2bp Ins | Frameshift | With above adjacent small INDEL reading frame restored. *Not linked to resistance.* |
| LOC110370195 | tRNA-splicing endonuclease subunit Sen34 | 1bp Ins | Frameshift | Potentially mis-called, only a single read supports insertion. SCD and HARM1.0 reference support this sequence variant. *Not linked to resistance.* |

Short-read structural variant detection

Four genes carried SVs intersecting CDS or UTR regions in at least one resistant individual. All candidate SVs were either inconsistently present across resistant individuals or were also present in susceptible genomes. No SV identified from short-read data matched the expected recessive inheritance pattern. Thus, short-read structural variant callers failed to identify any variant plausibly linked to resistance.

**Table S6. List of structural variants with some presence in resistant individuals identified using Smoove.**

| **Transcript identifier** | **Gene ID and Gene Name** | **Structural variant type** | **Structural variant length** | **Annotation type** | **Comment. *Linked to resistance?*** |
|---|---|---|---|---|---|
| rna-XM_021325770.1 | LOC110370060: tether containing UBX domain for GLUT4 | DEL | -163 | 3' UTR | Deletion consistent across HARM1.0 and SCD references. *Not linked to resistance.* |
| rna-XM_021325911.1 | LOC110370171: lipase 3-like | DEL | -5781 | CDS | Deletion only present in 1 individual as a heterozygous variant with little support, all other resistant individuals do not have this SV. *Not linked to resistance.* |
| rna-XM_021325929.2 | LOC110370188: uncharacterized transmembrane protein DDB_G0289901 | DEL | -114 | CDS | Highly variable gene between HaSCD and reference. Larger deletion seen in reference at this point. *Not linked to resistance.* |
| rna-XM_049850982.1 | LOC110370191: uncharacterized protein | DEL | -68 | 3' UTR | Deletion not present in all 17_294 $F_1$ individuals (missing in MIC $F_0$ Resist and FIC $F_1$ Resist). *Not linked to resistance.* |

Long-read structural variant detection

Sniffles identified 1,226 SVs in the resistance interval (654 deletions, 560 insertions, and 12 breakpoint/inversion events). Intersection with gene annotations and comparison across MIC individuals reduced these to a small number of candidates shared across all resistant individuals and absent in the susceptible parent.

**Table S7. List of structural variants with some presence in resistant individuals identified using Sniffles.**

| Locus ID | Gene description | SV type / size | Genomic location affected | Inheritance / validation outcome |
|---|---|---|---|---|
| LOC110370188 | Uncharacterized transmembrane protein | 160 bp insertion | Coding region | Present in resistant individuals but heterozygous in an additional 17-294 sample; not linked to resistance |
| LOC126056805 | Thyroglobulin-like | ~16 kb insertion with two adjacent “breakpoints” | Exon 5 | Homozygous in all resistant individuals and heterozygous in F1; insertion in exon leads to addition of 16 amino acids followed by a premature stop codon |
| LOC110370049 | Ecdysoneless homolog | Structural variant | 5′ UTR | Matches MIC inheritance pattern but failed validation in an additional resistant sample |

Large insertion investigation

The disruptive event identified from the Sniffles structural variant caller in the thyroglobulin-like gene were focused on a particular area of exon 5. In the short-read and long-read alignments at this area a distinct 9 base-pair 'bump' can be identified in the coverage of resistant individuals (Supplementary Figure S6). To characterise what is occurring at this position the region was identified in the long-read *de novo* assembly of the homozygous '17-294' individual. Mapping of the assembly again produced the characteristic 'bump' at this location where two sequences overlap (Supplementary Figure S6), and in this instance it is the same contig mapping over this region with a small overlap.

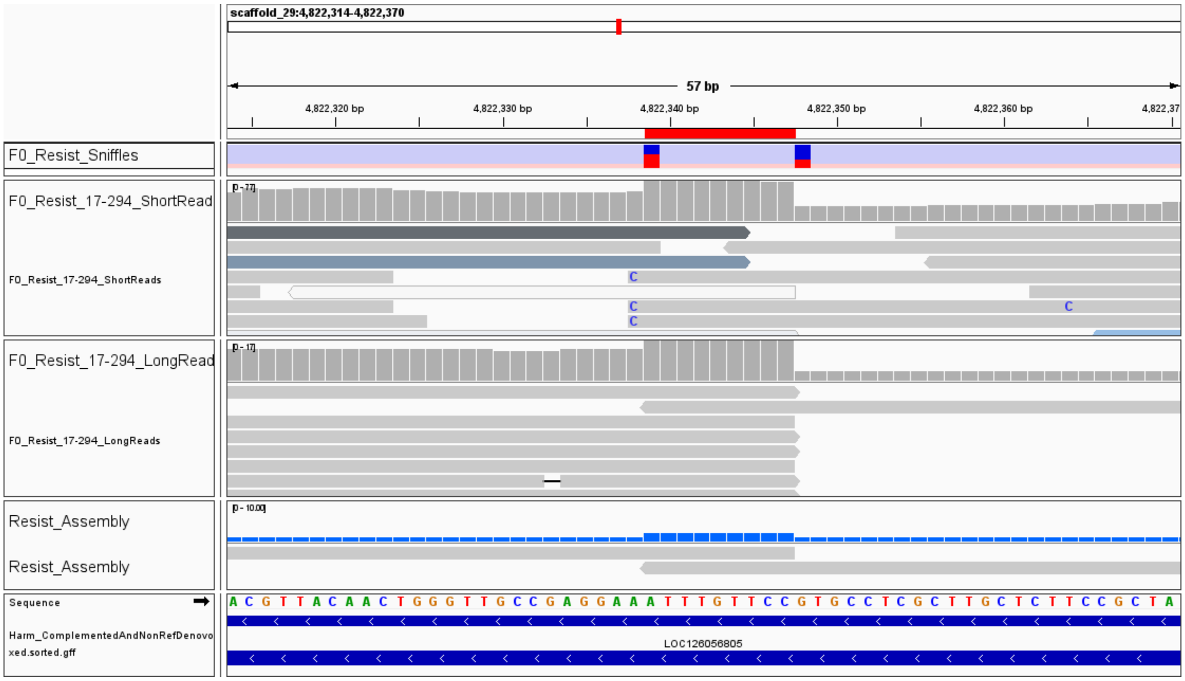


**Figure S6. Characteristic 'bump' in read coverage at the insertion site from mapping of resistant individuals to the susceptible reference.** The insertion site is present in the coding sequence of exon 5 of the thyroglobulin-like gene (LOC126056805). Short-read data was aligned using BWA-MEM2 and long-read data was aligned using minimap2. Alignments of short-reads have high clipping and missing paired end reads at this region, and alignments of long-reads have partial alignments to other regions after and before the insertion site. As described below, the action of the transposable element causes a Target-Site Duplication (TSD) in the reference sequence, causing 9bp to be copied and leading to partial overlap of the short and long-read sequence data.

The classification by Sniffles of this event as a "breakpoint" event was due to the partial mapping of long-reads to the thyroglobulin-like gene as well as regions on chromosome 20 and 22. At these regions a consistent pattern of two inverted repeat elements was observed, a 415bp element identified by RepeatModeller as 'rnd-6_family-1907#DNA'. Alignment of this element to the repeat region identified by Geneious in the insertion region shows 99% nucleotide identity. This points toward other potential sites of this transposable element in the genome, and the type of insertion as related to type II class DNA transposable elements.
